# Connectomic Analysis of Mitochondria in the Central Brain of *Drosophila*

**DOI:** 10.1101/2024.04.21.590464

**Authors:** Kit D. Longden, Patricia K. Rivlin, Michał Januszewski, Erika Neace, Louis K. Scheffer, Christopher Ordish, Jody Clements, Elliott Phillips, Natalie Smith, Satoko Takemura, Lowell Umayam, Claire Walsh, Emily A. Yakal, Timothy F. Musial, Michael J. Devine, Michael B. Reiser, Stephen M. Plaza, Stuart Berg

**Author notes:** **For correspondence:**; (KL, PR, SB). These authors contributed equally to this work (Kit D. Longden, Patricia K. Rivlin, Michał Januszewski, Erika Neace, Louis K. Scheffer).

## Abstract

Mitochondria are integral to the metabolism and cell biology of a neuron. Electron microscopy images of fly brain volumes, taken for connectomics, can be analyzed for mitochondria as well as the cells and synapses already reported. Here, from the *Drosophila* Hemibrain connectome dataset, we extract, classify, and measure approximately 6 million mitochondria, with the majority located among more than 20 thousand neurons and over 5500 cell types. Each mitochondrion is annotated with its location, orientation, voxel size, and appearance (dark and dense, light and sparse, or medium), and each synapse is linked to its closest mitochondrion. Using these data, we show how the most basic characteristics of mitochondria—volume, distance from synapses, and appearance—vary considerably between cell types and between brain regions. Mitochondria are larger and closer at presynapses than at postsynapses, and presynapses typically have a mitochondrion within one micron. However, cells important for learning and memory, Kenyon cells, have unusually small and sparse mitochondria that are placed with unusual precision near only half of presynapses. We find that mitochondria occupy a greater fraction of cell volume in inhibitory neurons than excitatory cells, dopaminergic neurons have a distinct synaptic positioning of mitochondria, and that glutamatergic neurons have a high fraction of mitochondria with a light appearance. We also find that synapses with more postsynaptic partners have larger presynaptic mitochondria and more distant postsynaptic mitochondria. Finally, we have extended the data model of our public web interface, neuPrint, to record our mitochondria data as searchable subcellular components of neurons in a connectomic database. These results indicate wide-ranging principles for how mitochondrial organization varies by cell type in a dataset of unprecedented size and coverage.

## Introduction

Mitochondria are cell organelles with many essential functions, including the regulation of intracellular calcium, cell signaling, the synthesis of various neurotransmitters, and the production of ATP, needed by the many energy-consuming processes in each cell. They are dynamic and are trafficked to sites of high energy demand (***Lewis and Lewis, 1915***; ***Boldogh and Pon, 2007***; ***Chang et al., 2006***; ***Schwarz, 2013***; ***Sheng, 2014***), including synapses, which have a high metabolic cost (***Karbowski, 2012***; ***Attwell and Laughlin, 2001***). While many synapses have nearby or adjacent mitochondria, many do not, and the relationship between synapses and mitochondria has been the subject of numerous studies (***Chavan et al., 2015***; ***Smith et al., 2016***; ***Schubert et al., 2022***; ***Vaccaro et al., 2017***; ***Sun et al., 2013***; ***Turner et al., 2022***; ***Devine and Kittler, 2018***). Here, we build upon recent advances in connectomics of the adult *Drosophila* brain to generate an analyzed and searchable dataset that contains an unprecedented number of mitochondria and neuronal cell types, and includes annotations of the distances between synapses and mitochondria.

The analysis of mitochondria from electron microscopy (EM) images has a long history (***Claude and Fullam, 1945***; ***Palade, 1953***). There are publicly available datasets of such images (***Lucchi et al., 2011***; ***Casser et al., 2020***), for example in 3D in mammals (***Wei et al., 2020***), 2D in a wide variety of organisms (***Conrad and Narayan, 2023***), and in pyramidal cells of mouse cortex (***Turner et al., 2022***). Volume EM images of the *Drosophila* brain offer the potential to investigate the synapse-mitochondria relationship in an additional organism, with high resolution and far greater numbers of synapses, cell types, and mitochondria.

For our analysis, we used EM images collected for the Hemibrain connectome, a sample comprising ∼2/3 of an adult *Drosophila* central brain, with >20,000 neurons, >5,500 cell types, and >18 million chemical synapses (***Scheffer et al., 2020***). Hemibrain data are accessible through our publicly available portal neuPrint (***Plaza et al., 2022***), where a segmented neuron is annotated as ‘traced’ if all of its main branches within the volume are reconstructed, ‘named’ if they have convincing identity information, as assessed by morphology, connectivity, and comparisons between hemispheres and with other cells, and ‘uncropped’ if most of its arbors are contained in the volume (excluding the soma; ***Scheffer et al. 2020***). Despite this definition of ‘uncropped’, many cells have substantial cropping of their processes, notably with cells with processes in the subesophageal zone, ventral nerve cord, and the optic lobes. Our analysis is focused on traced, named, and uncropped neurons, unless otherwise stated (Fig. 1A). In *Drosophila*, presynapses can be identified by the presence of a T-shaped bar structure (T-bar), and postsynapses can be identified by an electron dense thickening of the cell membrane opposing the presynapse, the postsynaptic density (PSD; ***Takemura et al. 2008***). The typical *Drosophila* synapse is polyadic, having multiple postsynaptic partners, and for this reason the number of presynapses is less than postsynapses (Fig. 1A).

**Figure 1.**
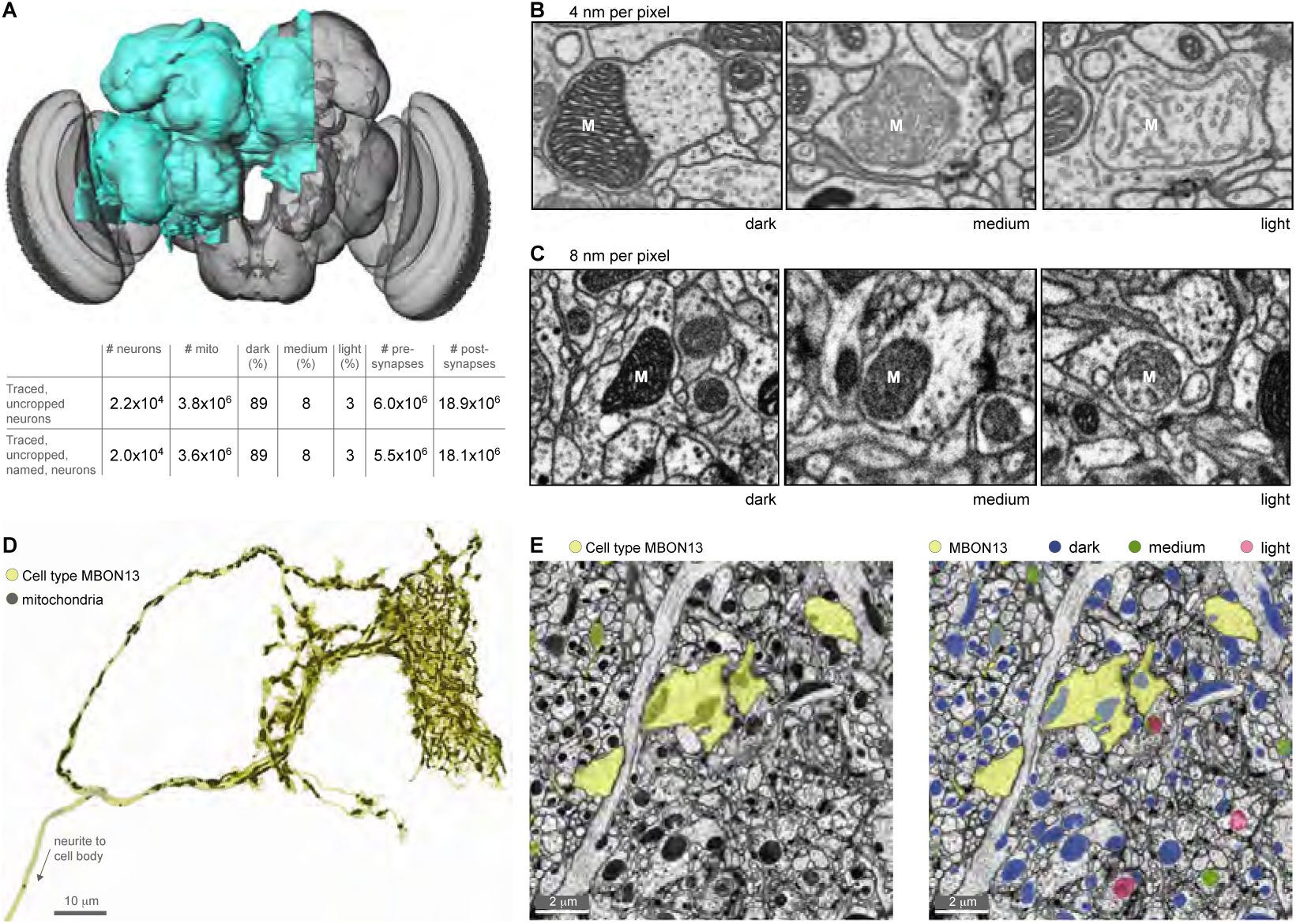
Mitochondria in the Hemibrain EM data. **A.** Grayscale rendering of the *Drosophila* brain, with the Hemibrain volume highlighted in cyan. Table lists numbers and properties of the dataset for neurons labeled in neuPrint: traced, uncropped and named neurons are the focus of this analysis. **B.** Images taken at 4 nm per pixel that illustrate differences between mitochondria appearance types in detail. Dark mitochondria are densely packed with lamellar cristae, have dark cytoplasmic density and are often elongated. Medium mitochondria have disorganized cristae and elongate-to-round shape. Light mitochondria have dilated cristae and an ovoid-to-round shape. **C.** Images from Hemibrain dataset at 8 nm per pixel, illustrative of the data analyzed. M denotes mitochondrion. **D.** Example of a segmented cell type, MBON13, rendered in yellow, and its mitochondria rendered in dark gray. **E.** View of this neuron, MBON13, in EM slices, segmented in yellow, and visualized without (left) and with (right) segmented mitochondria. Mitochondrial appearance types are labeled with blue for dark, green for medium, and cherry for light.

The Hemibrain images were taken using scanning electron microscopes with focused ion beam milling between sections and an isotropic resolution of 8 nm/voxel. Although the sample preparation was optimized for high-quality reconstruction of neurons and their synapses, mitochondria are well stained and preserved, enabling automated classification and segmentation at 8 nm/pixel (***Scheffer et al., 2020***), and 3D reconstruction to reveal mitochondrial internal structure at 4 nm/pixel (***Lu et al. (2022)***; Fig. 1B). Across species, organs, and preparations, mitochondria come in a wide variety of shapes, sizes, and structures (***Munn, 2014***). Following prior work that has used the appearance of mitochondria to classify cell types and axon terminals (***Delgado et al., 2019***; ***Klooster and Vrensen, 1997***; ***Li et al., 2003***), we used the appearance of mitochondria in our images to classify them as belonging to one of three appearance types: light, medium, or dark (Fig. 1B). It is not straightforward to link mitochondrial appearance types to those described in the literature, for example the ‘orthodox’ and ‘condensed’ configurations introduced by the seminal work of Hackenbrock (***Hackenbrock 1966***; see Methods for a discussion of this issue), and we report all our results in terms of our mitochondrial appearance types. Within our categories, dark mitochondria have tightly packed cristae (Fig. 1B), consistent with high ATP production, while light indicates more open cristae, indicating a greater role in biosynthesis. However, the physiology of mitochondria depends on variables without structural correlates in EM images, such as calcium concentration and membrane voltage, and therefore cannot be directly inferred.

The sample contains, in total, approximately 6 million mitochondria, with >3.5 million in traced, named, and uncropped neurons (Fig. 1A). To the knowledge of the authors, this is by far the largest analyzed dataset of mitochondria. Building on previous work classifying cell types from neuronal morphology, connectivity and genetic driver lines (***Scheffer et al., 2020***; ?; ***Schlegel et al., 2024***), we have been able to quantify the distributions of many different properties of mitochondria, including their volume, number, appearance, and distances from synapses, across an unprecedented variety of neuronal cell types. In particular, we have calculated for each synaptic element, both pre- and postsynaptic, the distance and size of the closest mitochondrion. These experimentally determined distributions are a significant contribution of this paper and have been added to our publicly available portal, neuPrint, where they can be searched, analyzed and visualized (***Plaza et al., 2022***).

## Results

### Mitochondrial properties and cell types

Using our segmentation of mitochondria in the Hemibrain dataset, we have quantified, for the first time, how mitochondria vary across thousands of identifiable, named neuronal cell types (Fig. 2). For each mitochondrial property examined, the distributions are unimodal, allowing us to identify cell types with unusually high or low values, for example, cell types with values lying beyond the 5^th^ and 95^th^ percentiles (Fig. 2B-L, Top). The variability within each cell type, measured by the standard deviation, is consistently small across all the different cell types, relative to the range of the mean values (Fig. 2B-L, Bottom). These data quantify the natural ranges of mitochondrial properties in *Drosophila* neurons, which were not previously known, and demonstrate that the variability within cell types can be sufficiently low to identify cell types with unusual values.

**Figure 2.**
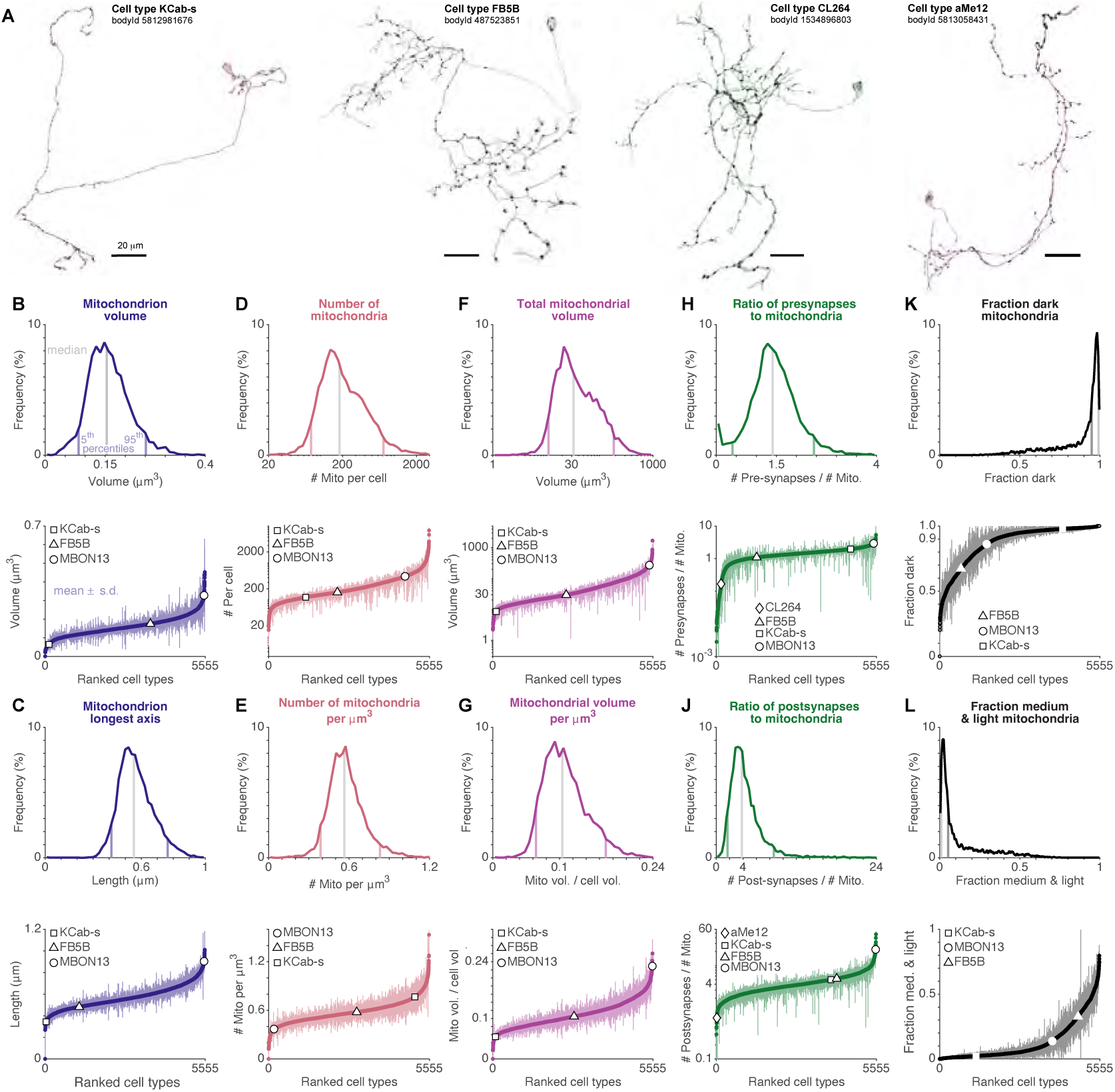
Mitochondrial properties of cell types, with unusual examples including cell types involved in learning and memory. **A.** Examples of segmented cell types and their mitochondria, chosen to collectively illustrate the ranges of mitochondrial properties. Symbols corresponding to these cell types, along with cell type MBON13 shown in Fig. 1D, are indicated in the lower plots of panels B-K; bodyId values identify them in neuPrint. Scale bars: 20 µm. **B.** Top: distribution of the mean median mitochondrion volume of each cell type; gray line indicates distribution median and vertical color lines indicate the 5th and 95th percentiles. Bottom: mean ±standard deviation of the median mitochondrion volume for every cell type, with the cell types ranked by the mean median value. The example cell types from panel A show that KCab-s cells have a small average mitochondrion volume, while MBON13 cells have large average mitochondrion volume, relative to other cell types. Panels **C-L** are organized as for B, but showing data for: **C** Longest axis of an ellipsoid fitted to each mitochondrion; **D** Number of mitochondria per instance of the cell type; **E** Number of mitochondria divided by the volume of the cell, for every instance of the cell type; **F** Total mitochondrial volume per instance of the cell type; **G** Total mitochondrial volume divided by the cell volume, for every instance of the cell type; **H** Number of presynapses divided by the number of mitochondria, for every instance of the cell type - some cells have no presynapses, and these are not shown; **J** Number of postsynapses divided by the number of mitochondria, for every instance of the cell type; **K** Fraction of dark mitochondria; and L Fraction of medium and light mitochondria combined.

To illustrate how individual cell types vary in their mitochondrial properties, we selected examples that span the observed ranges (Fig. 2A, 1D). Kenyon Cells (KCs) play an important role in the encoding of odors, learning and memory, and their mitochondria contribute to the regulation of synaptic plasticity (***Comyn et al., 2024***; ***Pavlowsky et al., 2024***). An example of a KC type, a KCab-s cell, is shown in Fig.2A (left), rendered with its mitochondria. The mitochondrial properties of all KCab-s cells (n=223 cells) are indicated by square symbols across Fig. 2B-L. Their mitochondria are small (Fig. 2B,C; percentiles: vol. 2.4%; len. 0.5%), few in absolute number (Fig. 2D; percentile: 23%), but many relative to their slender cell volume (Fig. 2E; percentile 91%). They occupy a small volume of the cell (Fig. 2F,G; percentiles: total vol. 2.0%; vol. dens. 1.6%), have a typical-to-high ratio of synapses to mitochondria (Fig. 2H,J; percentiles: ratio pre. 84%; ratio post. 72%), and a high proportion of dark mitochondria, indicating compact cristae structure (Fig. 2K,L; percentiles: dark 77%; med. & light 23%). These properties are typical of all KC cell types (Suppl. Data Tables 1,2), and show that while their synaptic plasticity is critically dependent on the energy supply (***Plaçais et al., 2017***), this is achieved with a low investment of small mitochondria per cell, consistent with the sparse and low activity levels observed in KCs (Honegger et al., 2011).

In contrast to KCs, their downstream cell types can be highly invested with mitochondria. As an example, the mushroom body output neuron (MBON) cell type MBON13 is downstream of KCa’b’ cells, and is shown with its mitochondria in Fig. 1D. This cell type is specifically required for a complex form of learning, second-order conditioning, where an association is learned between a new odor and an already conditioned odor (***Rachad et al., 2025***). The mitochondrial properties of the MBON13 cell (n=1 cell) are indicated by circle symbols across Fig. 2B-L. The MBON13 mitochondria are large (Fig. 2B,C; percentiles: vol. 99.6%; len. 99.5%), many in absolute number (Fig. 2D; percentile 85%), but few relative to the cell volume of their profuse processes (Fig. 2E; percentile: 3.3%). They occupy a large volume of the cell (Fig. 2F,G; percentiles: total vol. 98%; vol. dens. 99.9%), and have a high ratio of synapses to mitochondria, particularly the postsynapses that receive the inputs to the cells (Fig. 2H,J; percentiles: ratio pre. 98.6%; ratio post. 99.8%). They also have an average proportion of dark mitochondria (Fig. 2K,L; percentiles: dark 29.4%; med. & light 70.5%). This is a cell type that requires great mitochondrial investment: MBON13 has the highest density of mitochondria by volume of all MBON cell types, which themselves have twice the mitochondrial density of KC cell types (MBONs 14.5 ±3.4%, KCs 7.3 ±1.4%).

Specific cell types can have mitochondria that differ in a single property. For example, FB5B (Fig. 2A, second from left) is found in the fan-shaped body (FB), a brain region variously associated with locomotion, sleep and arousal (***Keleş et al., 2025***; ***Sarnataro et al., 2025***; ***Ni et al., 2019***), along with diverse sensory processing, including for navigation (***Currier et al., 2020***; ***Weir et al., 2014***; ***Hu et al., 2018***; ***Lyu et al., 2022***), and is a cell type that has broadly typical mitochondrial properties (Fig. 2B-J, triangle symbols, n=6 cells). However, it is unusual in having a relatively high proportion of mitochondria with a light or medium appearance type (Fig. 2K,L; percentiles: dark 13%; med. & light 87%). The majority of cell types contain mainly dark mitochondria, with a few cell types reaching 50% light or medium (Fig. 2K,L).

For mitochondrial properties relating to synapses, it is important to consider the limitations of the sample, which truncates many neurons particularly those with processes in the optic lobes and the subesophageal zone (SEZ). For example, cell type CL264 (Fig. 2A, second from right), named for its processes in the clamp brain region, has axons in the SEZ and outside the Hemibrain volume (***Berg et al., 2025***), and a correspondingly low ratio of presynapses to mitochondria in the Hemi-brain (Fig. 2H; percentile: 2.9%). Meanwhile, cell type aMe12 (Fig. 2A, right), a cell type that plays a key role in determining the color circuitry in optic lobe connectomes (***Kind et al., 2021***; ***Nern et al., 2025***), receives many inputs in the optic lobes and few in the central brain (***Nern et al., 2025***), and therefore has a low ratio of postsynapses to mitochondria in the Hemibrain (Fig. 2J; percentile: 0.3%). Given these limitations, the data show that synapses outnumber mitochondria across the many and varied cell types, such that few cells have a mitochondrion for every synapse.

Overall, these data quantify the ranges of mitochondrial properties across cell types, establish that there are consistent cell type differences, and show that individual cell types can differ from typical values in one or many of their mitochondrial properties.

### Mitochondrial properties and brain regions

Recent work in human tissue has indicated that mitochondrial density may vary by brain region (***Mosharov et al., 2025***). We therefore explored this issue in our data, benefiting from our greater spatial resolution (isotropic 8 nm voxels, compared to isotropic 3 mm voxels in one coronal slice in ***Mosharov et al. 2025***). By visual inspection, mitochondrial properties vary across the sample. The analysis of this variability is affected by differences in the level of reconstruction between areas: brain regions involved in olfaction, navigation, learning and memory, including the mushroom body (MB) and central complex (CX; Fig. 3A), are reconstructed to a higher level of completion than other brain regions (***Scheffer et al., 2020***). Therefore, our analysis is focused on MB and CX, and for clarity and robustness, we have grouped individual brain regions of interest (ROIs) into super ROIs (SROIs), following the naming conventions of ***Ito et al. (2014)***.

**Figure 3.**
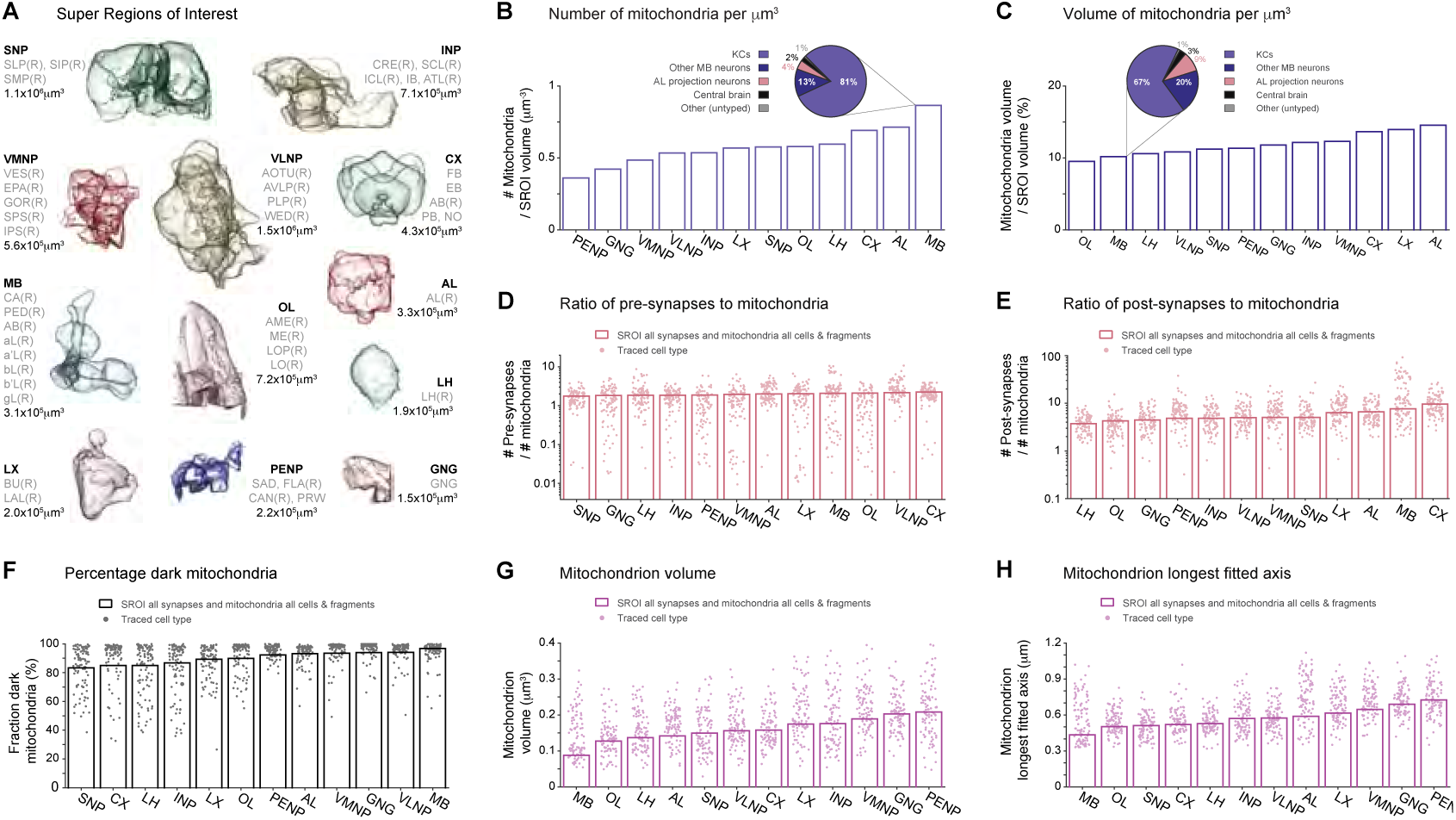
Mitochondrial properties vary by brain region, and vary more by cell type. **A.** Renderings of Hemibrain brain regions, organized as super regions of interest (SROIs), with lists of their constituent ROIs and volumes. SROIs are presented to give a visual appreciation of relative sizes; for their relative positions in the brain see ***Scheffer et al. (2020)***. SROIs are calculated for the right hemisphere (R) because the left hemisphere contains many truncated ROIs. **B-H**: Bars indicate values calculated using all cells, including cell fragments. Dots indicate values of traced, uncropped, named cell types, that make the greatest contribution to the number of mitochondria in the SROI, ranking ≤100. **B,C.** Numbers of mitochondria per unit volume (B) and percentage of SROI volume occupied by mitochondria (C). Insets: pie charts of how KCs and other groupings of neurons contribute to the MB total; MB local interneuron cell types, MBON, PAM, and PPL cell types comprise ‘other MB neurons’. **D,E.** Ratio of the number of presynapses (D) and postsynapses (E) to mitochondria in the SROIs. **F.** Percentage fraction of mitochondria with dark appearance type. **G-H.** Median mitochondrion volume (G) and longest axis of an ellipse fitted to each mitochondrion (H).

The mushroom bodies have the greatest number of mitochondria per unit volume (Fig. 3B). There is substantial variability between brain regions, with the number of mitochondria per unit volume varying by a factor of 2 between the lowest and highest values (Fig. 3B; PENP 0.36 µm^-3^; MB 0.87 µm^-3^). Kenyon cells contribute the majority of MB mitochondria (Fig. 3B; 81%, inset) and consistent with their low mitochondrial density, the MB mitochondrial volume density is low (Fig. 3C; MB 10.2%). Across all SROIs, the volume density varies by 50% (Fig. 3C; OL 9.5%, AL 14.5%). In the brain regions along the well-reconstructed olfactory pathway, the mitochondrial investment is greatest in the antennal lobes, where peripheral sensory input is processed, and then lower in the downstream mushroom bodies and lateral horn (Fig. 3C; LH 10.6%).

The central complex has the highest ratio of synapses to mitochondria (Fig. 3D,E). The ratio of presynapses to mitochondria is fairly constant across brain regions at just under two synapses per mitochondrion, varying by 28% between lowest and highest values (Fig. 3D; SNP 1.8, CX 2.2). This consistency is remarkable given the wide variability in the median values for cell types that contribute the most mitochondria to each SROI (Fig. 3D; 1st, 3rd quartiles for displayed cell types: CX 1.5, 2.9; MB 0.6, 2.9). Compared to the numbers of presynapses, the numbers of postsynapses per mitochondria are greater and vary more widely by brain region (Fig. 3E; LH 3.7, CX 9.6). Again, there is wide variability in the median values for cell types that contribute the most mitochondria to each SROI (Fig. 3E; 1st, 3rd quartiles for displayed cell types: CX 6.5, 12.4; MB 5.6, 24.8).

The mushroom bodies and central complex differ in their mitochondrial appearance types: the fraction of dark mitochondria is highest in MB, while the complementary fraction of medium and light mitochondria is high in CX (Fig. 3F; fraction dark: MB 97%; CX 85%; fraction medium and light: MB 3%; CX 15%). MB mitochondria also have the smallest volume and shortest fitted axis of all the brain regions we analyzed (Fig. 3G,H; *p* < 0.001, one-way ANOVA with Bonferroni correction), while mitochondria in CX and all brain regions show a high variability in these mitochondrial properties (Fig. 3G,H; 1st, 3rd quartiles for displayed cell types: volume MB 0.09, 0.19 µm^3^; CX 0.13, 0.19 µm^3^; longest axis MB 0.39, 0.71 µm; CX 0.48, 0.61 µm).

Mitochondrial properties can also vary between brain regions within an individual cell type. For example, in cell type CL042, 87% of the mitochondria are dark in the SNP brain region, but only 29% of mitochondria are dark in the SIP brain region (CL042, SNP n_mito_ = 147; SIP n_mito_ = 125). However, the median fractions of dark mitochondria of all cell types with ≥50 mitochondria in these two brain regions are strikingly similar (INP 94.4%, SNP 94.6%), and the presence of a cell process in an SROI does not strongly determine the local mitochondrial properties in that SROI.

Together, these data indicate substantial differences in mitochondrial distribution and properties between well-reconstructed brain regions. They also demonstrate a consistently greater variability in mitochondrial properties between cell types than between brain areas.

### Distances and sizes of mitochondria near synapses

Synaptic transmission carries a major energy cost for neurons, and mitochondria can locate close to active synapses where their energy is needed (***Harris et al., 2012***; ***Smith et al., 2016***; ***Zhu et al., 2021***). The release of synaptic vesicles at the presynapse is also critically dependent on the regulation of local intracellular calcium concentrations, which can be affected by the presence or absence of a nearby mitochondrion (***Kwon et al., 2016***; ***Vaccaro et al., 2017***; ***Li et al., 2020***; ***Hirabayashi et al., 2017***). We therefore analyzed whether different cell types would have different relationships in the positions and sizes of their mitochondria relative to their synapses (***Schubert et al., 2022***; ***Cserép et al., 2018***; ***Justs et al., 2022***; ***Smith et al., 2016***; ***Schneider-Mizell et al., 2016***). In the Hemibrain dataset, synapse locations are represented by a single point, the result of an automatic synapse detection process trained on manually annotated centers of T-bars and PSDs (***Scheffer et al., 2020***; ***Huang et al., 2018***). For each cell, we calculated the internal distance to the nearest mitochondrion from every presynapse, and from every postsynapse. The distances were calculated from the point representing the synapse to the nearest edge of the nearest mitochondrion, along the shortest 3D path inside the cell, and are reported in neuPrint.

To illustrate differences in synapse-to-mitochondria distances in individual cells, we provide three examples in Fig. 4. In the example KCab-c cell (Fig. 4A-F), only half the presynapses in the right alpha lobe brain region, aL(R), have a mitochondrion < 1 µm (Fig. 4A). The narrow axons are thicker around the mitochondria, and many en passant synapses lack nearby mitochondria (Fig. 4D-F). In contrast, the example FB3E cell (Fig. 4G-L) lacks en passant synapses in the fan-shaped body (FB) brain region, with presynapses occurring at varicosities that usually, although not always, contain a mitochondrion (Fig. 4J-L). Finally, the example FB5E cell (Fig. 4M-Q) has, like the majority of KCs, a low fraction of presynapses with a mitochondrion < 1 µm (Fig. 4M). However, in this cell type, many presynapses without a nearby mitochondrion are found at varicosities (Fig. 4O-Q). For all three cells, mitochondria are relatively sparsely distributed in connecting neurites, a common feature of many cell types.

**Figure 4.**
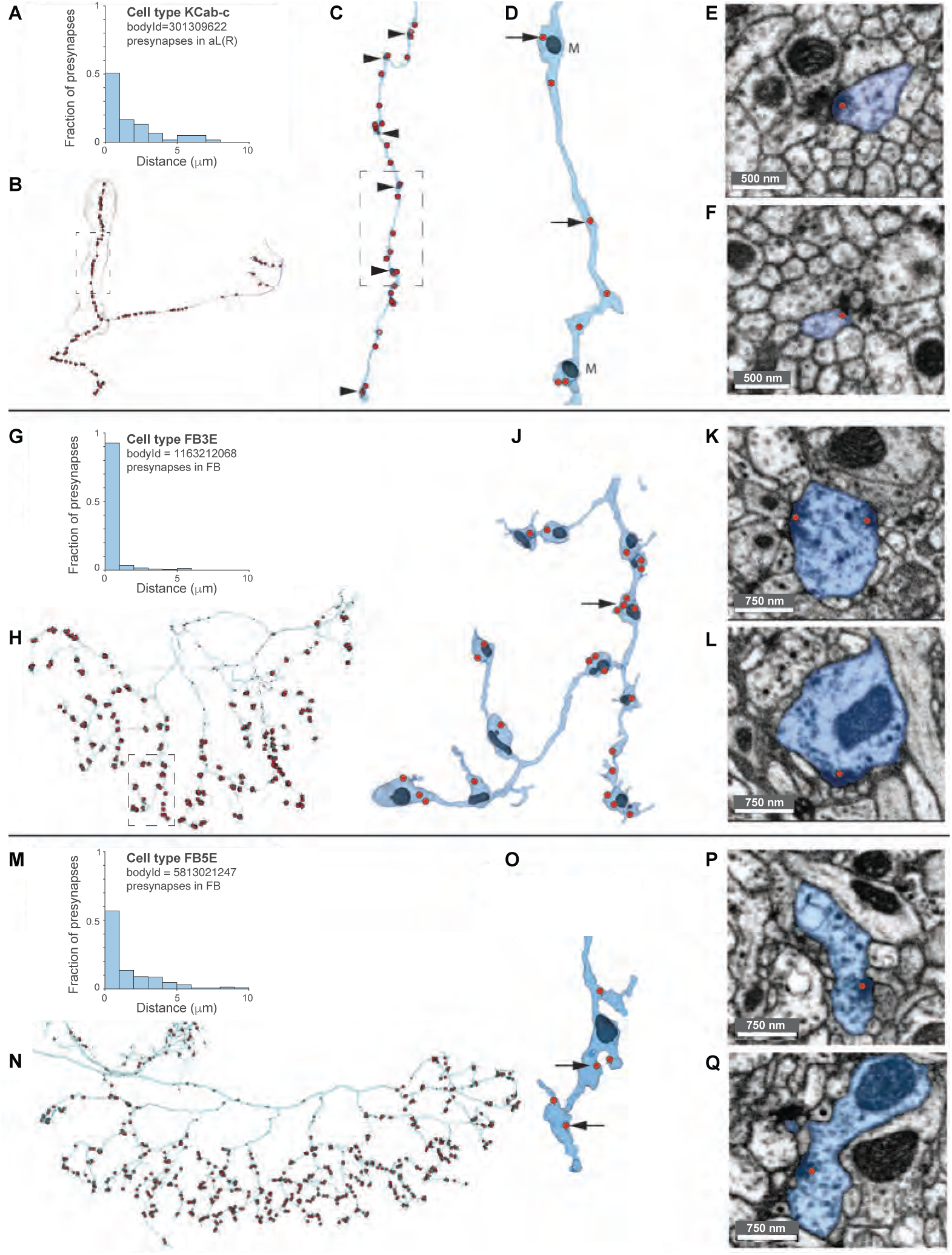
Spatial arrangements of presynapses and mitochondria. Example KCab-c cell (**A-F**), FB3E cell (**G-L**), and FB5E cell (**M-Q**). **A.** Fractions of presynapses with nearest mitochondrion distance in µm: only half presynapses have a mitochondrion < 1 µm. **B.** Reconstructed neuron, with presynapses (red dots), mitochondria (black), and right alpha lobe (aL(R)) brain region (gray). Dashed box indicates region of enlarged view in panel C. **C.** Enlarged view of the cell with mitochondria (black triangles). **D.** Further enlarged neuron section. Arrows indicate locations of EM views in panels E and F. **E,F.** EM views of presynapses (red dots); cell is labeled in blue. **G.** More than 90% of this FB3E cell’s presynapses in FB have a mitochondrion < 1 µm. **H.** Cell’s FB neurites. Dashed box indicates region of enlarged view in panel J. **J-L.** Enlarged view of neurites. Arrow indicates location of EM views in panels **K** and **L. M.** Less than 60% of this FB5E cell’s presynapses in FB have a mitochondrion < 1 µm. **N.** Cell’s FB neurites. **O-Q.** Enlarged view of a neurite, with viewing angle altered from panel M. Arrows indicate locations of EM views of presynapses in panels **P** and **Q.**

To illustrate how consistently synapse-to-mitochondrion distances are organized within groups of functionally related cell types, we focus on three groups of cell types of the mushroom body (Fig. 5): KCs, which encode odors, and implement synaptic plasticity supporting olfactory learning; protocerebral anterior medial (PAM) cells, which provide neuromodulatory input required for learning; and MBONs, whose activities are correlated with behavioral attraction or repulsion. Even within this single brain region, different functional groups of cell types have distinct spatial arrangements of their synaptic mitochondria (Fig. 5A). For MBONs, the presynapses are close to mitochondria, indicative of relatively active outputs and tightly mitochondrially controlled intracellular calcium, while the postsynapses (inputs) span a wide range, from relatively close (MBON13) to far (MBON15; Fig. 5A). It is possible that some MBONs get relatively constant input, and others receive intermittently active or spatially sparse inputs, and these factors may account for the wide range. For KCs, the postsynapses are closer to mitochondria, compared to the other MB cell types, indicating their inputs may be more commonly activated (Fig. 5A). Meanwhile, their presynapses, which are a locus of synaptic plasticity, cover a wide range, consistent with differences in input activity levels and observed variability in presynaptic calcium levels (***Davidson et al., 2023***). For PAM cell types, the presynapses are further from mitochondria, relative to MBONs and most KCs (Fig. 5A), indicating a reduced mitochondrial requirement for their synaptic outputs than MBONs and KCs, on average.

**Figure 5.**
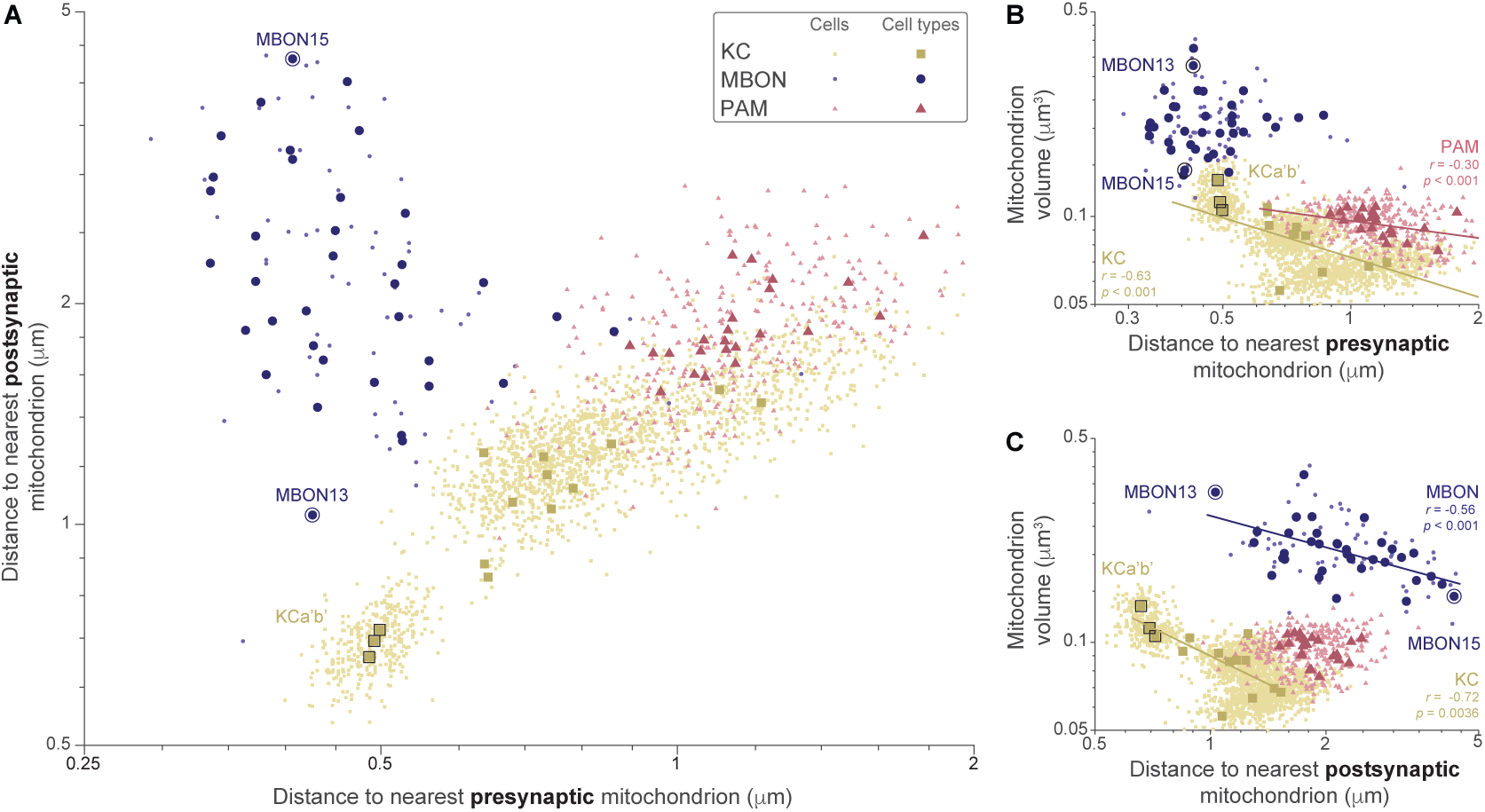
Consistent synapse-to-mitochondrion distances of MB cell type groups. Data for three MB cell type groups are shown: MBONs (blue circles), KCs (yellow squares), and PAMs (red triangles); each small symbol is a single cell, and each larger symbol is the median of a single cell type. In addition, two extremal MBON cell types are labeled with black symbol outlines, MBON13 and MBON15, as are extremal KCa’b’ cell types. **A.** Distances from pre- and postsynapses to the nearest mitochondrion. **B.** Mitochondrion volume versus distance from presynapses to the nearest mitochondrion. Lines indicate linear fits in the plotted log-log space (corresponding to power law fits in linear space) of the median values for individual cells, for groups of cells with significant correlations; fits are labeled with Pearson’s correlation coefficient (*r*) on log transformed data and the *t* statistic (*p* value). Fit for MBONs is not significant (*p* = 0.43). **C.** Mitochondrion volume versus distance from postsynapses to the nearest mitochondrion, with linear fits shown following conventions of panel B.

Since larger mitochondria may be expected to be closer to synapses, we investigated the relationship between the volume of the closest mitochondrion and its distance from a synapse (Fig. 5B,C). KC presynapses with smaller median distances to mitochondria have larger nearest mitochondria, an effect ranging over more than an order of magnitude in both mitochondrial volume and distance (Fig. 5B; KC, Pearson’s correlation coefficient on log transformed data, *r* = −0.74, *t* statistic *p* < 0.001). This relationship also holds for PAM cells, but not for MBONs (PAM *r* = −0.30, *p* < 0.001; MBON *p* = 0.43). In contrast, at postsynapses the relationship between the volume and distance of the closest mitochondrion varies considerably between cell type groups (Fig. 5C). For KCs and MBONs, the mitochondrial volume is inversely correlated with the distance, which may reflect the energy demands of the postsynapses of these cell types (Fig. 5C; KC, Pearson’s correlation coefficient on log transformed data, *r* = −0.72, *t* statistic *p* = 0.0036; MBON, Pearson’s *r* = −0.56, *t* statistic *p* < 0.001), but across all MB cells analyzed, mitochondrial volume weakly increases with distance (*r* = 0.29, *t* statistic *p* = 0.0117).

An important finding in rat hippocampal CA1 axons is that ≤ 50% of presynapses in synaptic boutons have a mitochondrion within 1 µm (***Smith et al., 2016***). In *Drosophila*, the axons of KCs are comparable in shape and size, and share other similarities, for instance both contain en passant synapses and are important sites of synaptic plasticity. Therefore, to facilitate comparisons with the analysis of ***Smith et al. (2016)***, we computed for every cell the percentage of pre- or postsynapses whose nearest mitochondrion was <1 µm, 1-3 µm, or >3 µm (Fig. 6). Across cells grouped by cell type, the majority of presynapses have a mitochondrion < 1 µm (Fig. 6A); the median fraction of presynapses with a mitochondrion < 1 µm is less than 65% for only 5% of cell types (Fig. 6B). KCs have low median cell type fractions, ranging 46-79%, with only KCa’b’ and KCg-s3 cell types having median fractions greater than the 5^th^ percentile of cell types (Fig. 6C). PAMs also have atypically low median cell types fractions, ranging 36-55% (Fig. 6C). In contrast, MBONs have median cell type fractions that are typical of all cell types, spanning the full range (Fig. 6A-C). For postsynapses, the fractions with mitochondria < 1 µm and > 3 µm are typically comparable across all cell types (Fig. 6D). KCs have unusually high fractions of postsynapses with a mitochondrion < 1 µm, ranging 39-65%, and 8 KC cell types having a median cell type fraction greater than the 95^th^ percentile (Fig. 6E,F). In contrast, PAMs and MBONs have median cell type fractions spanning the range of values for all cell types (Fig. 6F). Together, these results demonstrate that mitochondria are unusually located in KCs, with mitochondria far from presynapses and close to postsynapses, compared to all cell types examined. They also confirm that, like CA1 synapses in the rat hippocampus (***Smith et al., 2016***), not all presynapses involved in synaptic plasticity have access to nearby mitochondria. Collectively, these results indicate that mitochondria are located differently at presynapses and postsynapses. Cell types such as APL, one of the largest in the dataset, have tens of thousands of synapses, enabling the analysis of synapse-to-mitochondria distances within a single cell (Fig. 7A). Functionally, APL provides non-spiking inhibitory feedback to Kenyon cells in the mushroom body (***Amin et al., 2020***). In the APL cell, the distance to a mitochondrion is smaller for presynapses, on average, than for postsynapses (Fig. 7B; *p* < 0.001, Wilcoxon rank sum test on log transformed data). This relationship is typical: for 96% of cell types, the median distance to the nearest mitochondrion is lower for presynapses than for postsynapses (Fig. 7C). Overall, the distribution of nearest mitochondrion distances for presynapses and postsynapses is well covered by the mushroom body groups of cell types presented in Fig. 5, the PAM, KC and MBON cell type groups (Fig. 7D; PAM cherry triangles, KC yellow squares, MBON blue circles). For example, the cell types of the posterior slope brain region, the PS cell types, have values that collectively span the range (Fig. 7D; PS, green inverted triangles), and individual cell types account for outliers. For example, the population of olfactory receptor neurons ORN_DM6 (n = 60), have mitochondria unusually near to both presynapses and postsynapses (Fig. 7D; white diamond).

**Figure 6.**
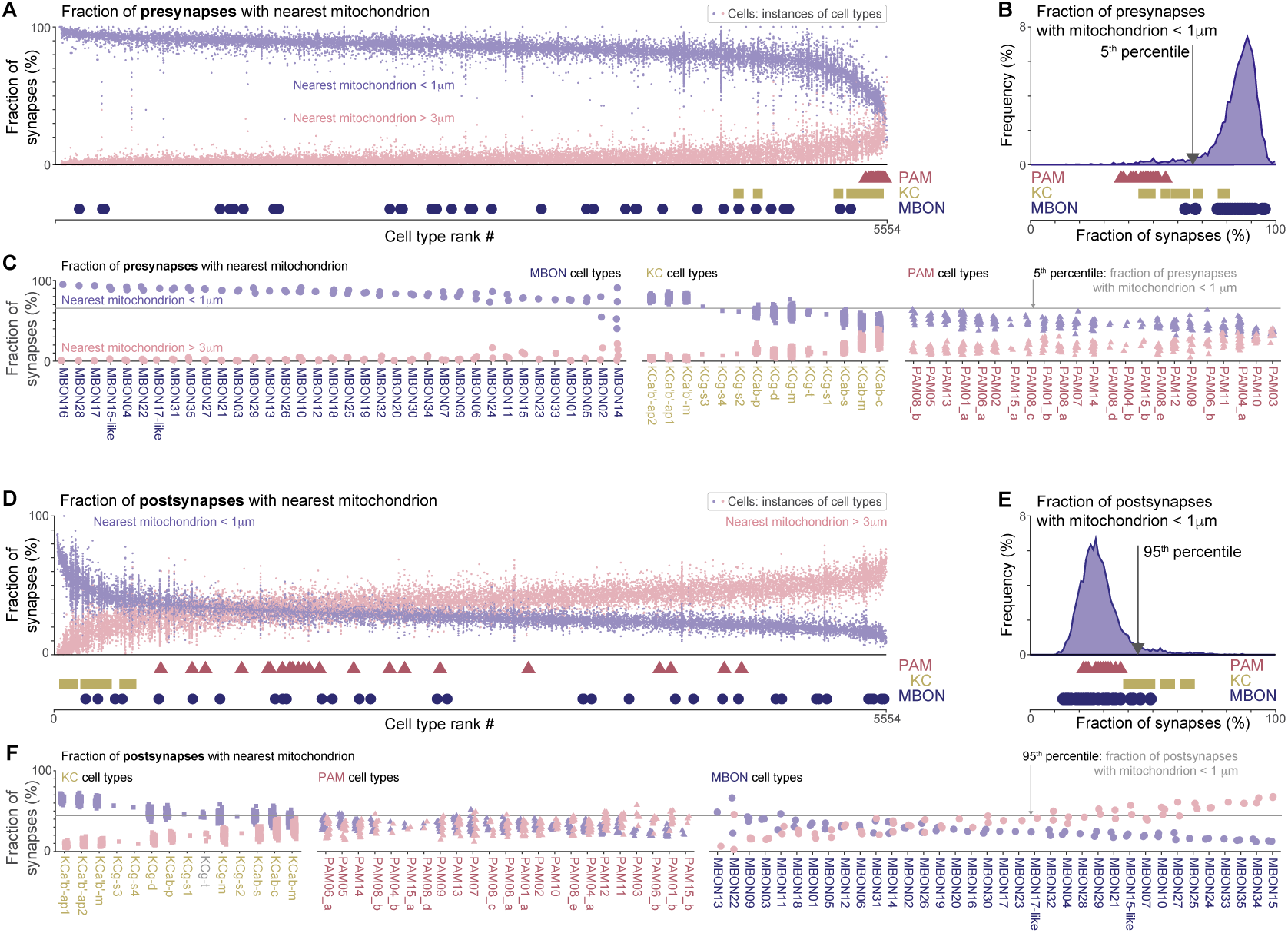
Majority of presynapses and minority of postsynapses have a nearby (< 1 µm) mitochondrion. **A.** Fraction of presynapses with mitochondria near (< 1 µm, blue) and distant (> 3 µm, pale cherry) for cells, instances of cell types (dot symbols). Cell types are ranked by median fraction of presynapses with mitochondria < 1 µm. Values of cell types from three cell type groups are indicated below the horizontal axis: PAMs (triangles), KCs (squares) and MBONS (circles). **B.** Frequency distribution of median cell type fraction of presynapses with mitochondria < 1 µm. Vertical line indicates 5th percentile value of 65.4%. Symbols below the horizontal axis indicate PAM, KC and MBON cell types. **C.** Cell type grouped data for, left-to-right, MBON, KC and PAM cell types, showing fraction of presynapses with mitochondria < 1 µm (blue) and > 3 µm (pink) for cells (dots). Within each cell type group (MBON, KC, PAM), cell types are ranked by median fraction of presynapses with mitochondria < 1 µm. Horizontal line indicates the 5th percentile value of 65.4%. **D.** Fraction of postsynapses with mitochondria < 1 µm and > 3 µm for cells, with plotting conventions as in panel A. **E.** Frequency distribution of median cell type fraction of postsynapses with mitochondria < 1 µm. Vertical line indicates 95th percentile value of 44.2%. **F.** Cell type grouped data for, left-to-right, KC, PAM, and MBON cell types, showing fraction of postsynapses with mitochondria < 1 µm and > 3 µm for cells. Horizontal line indicates the 95^th^ percentile value of 44.2%; other plotting conventions as in panel C.

**Figure 7.**
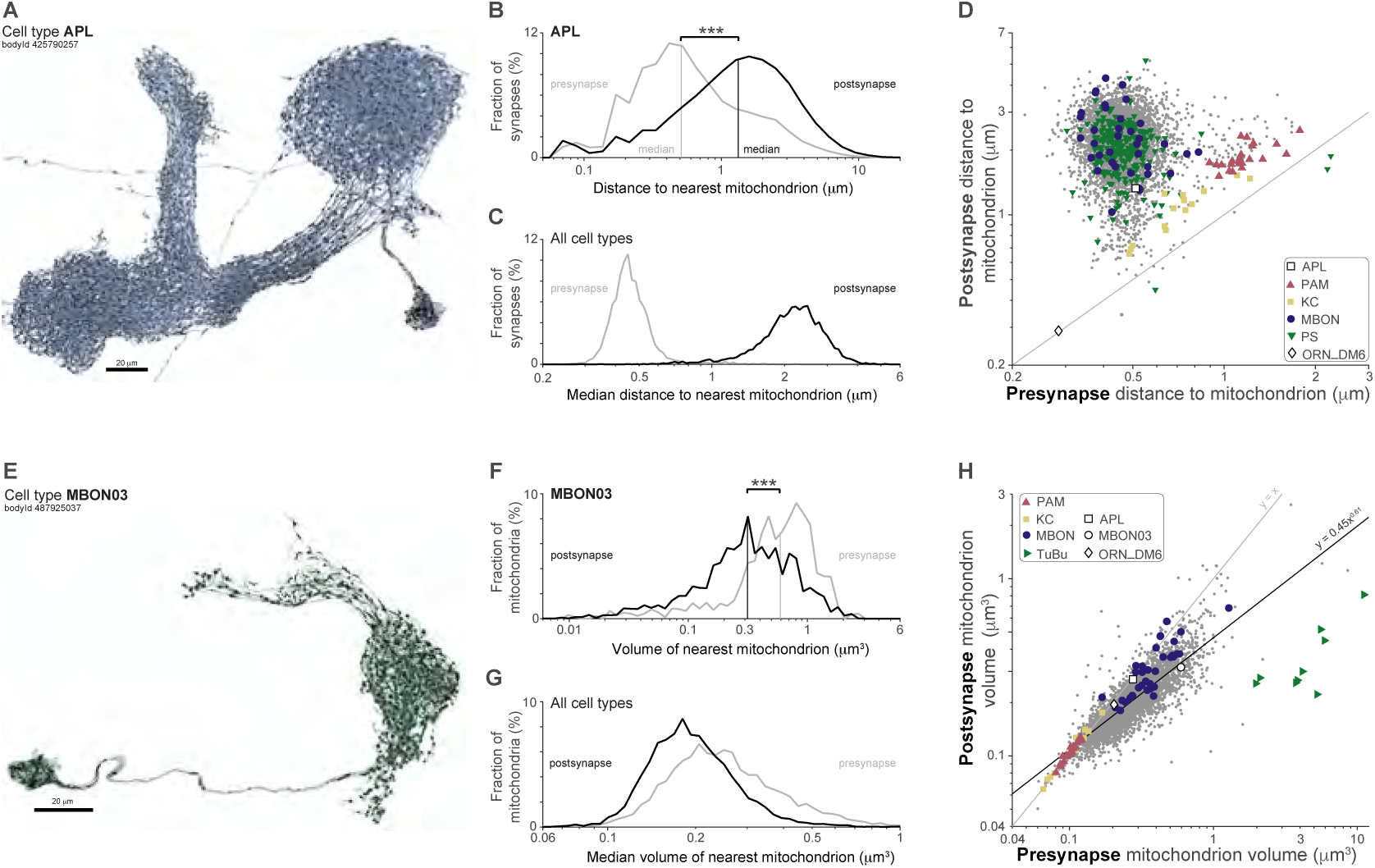
Mitochondria are closer and larger at presynapses than at postsynapses. **A.** Segmented APL cell (blue) and mitochondria (black). **B.** APL distributions of nearest mitochondrion distance for presynapses (gray) and postsynapses (black). Asterisks indicate significance: *** *p* < 0.001, Wilcoxon rank sum test on log transformed data. **C.** Distribution of all cell types median nearest mitochondrion distance for presynapses (gray) and postsynapses (black). **D** Joint distribution of median nearest mitochondrion distance for presynapses (horizontal axis) and postsynapses (vertical axis), for APL, PAM, KC, MBON, posterior slope (PS), ORN_Dm6 cell types (see legend for symbols), and all other cell types (gray dots). Cell type median values calculated from cells with >50 mitochondria, ≥50 presynapses and ≥50 postsynapses. Gray line indicates equal values. **E.** Segmented MBON03 cell (green) and mitochondria (black). **F.** MBON03 distributions of nearest mitochondrion volume for presynapses (gray) and postsynapses (black). Plotting conventions as panel (B). **G.** Distribution of all cell types median nearest mitochondrion volume for presynapses (gray) and postsynapses (black). **H** Joint distribution of median nearest mitochondrion volume for presynapses (horizontal axis) and postsynapses (vertical axis), for PAM, KC, MBON, neurons connecting the tubercle and bulb brain regions (TuBu), APL, MBON03, ORN_Dm6 cell types (see legend for symbols), and all other cell types (gray dots). Cell type median values calculated from cells with >50 mitochondria, ≥50 presynapses and ≥50 postsynapses. Dark gray line indicates linear best fit of logarithmically scaled values, equivalent to a power law relationship; pale gray line indicates equal values.

If mitochondria are closer to presynapses than to postsynapses, are they also larger at presynapses? In the APL cell type, the mitochondrial volume distributions are very nearly indistinguishable for the mitochondria closest to presynapses and postsynapses. However, in other cell types, the mitochondrial partners of presynapses and postsynapses may have different volumes, as illustrated by the MBON03 cell type (Fig. 7E). In this cell type the nearest mitochondria are larger at presynapses than at postsynapses (Fig. 7F), a trend that holds for all cell types (Fig. 7G). For 97% of cell types, the median volume of the nearest mitochondria is greater for presynapses than for postsynapses (Fig. 7H). Across all cell types, there is a trend for the nearest mitochondrion size to be correlated at presynapses and postsynapses, and is well described by linear fit in logarithmic space, corresponding to a power law relationship (Fig. 7H). Of the outliers to this trend, the neurons connecting the tubercle and bulb brain regions, the TuBu cells, are an intriguing group of cell types that collectively have unusually large nearest mitochondria to presynapses (Fig. 7H; TuBu green lateral triangles).

Together, the results of this section show that cell types, and groups of cell types, can have distinct patterns of synapse-to-mitochondrion distances (Fig. 5A) and that most presynapses have a nearby mitochondrion (Fig. 6A). Meanwhile, mitochondria are typically closer and larger at presynapses than at postsynapses (Fig. 7D,H). The KC and PAM groups of mushroom body cell types in particular have unusual placements of mitochondria near both pre- and postsynapses (Fig. 6), illustrative of the variety of spatial organization of mitochondria at synapses across cell types (Figs. 4-7).

### Stereotypy of synapse-to-mitochondrion distances within cell types and cells

Cell types differ in the stereotypy of synapse-to-mitochondrion distances, a variability that we could quantify in our dataset by analyzing the cell types with many (≥ 30) cells with many (> 50) mitochondria (Fig. 8). As before, we have focused on the well-reconstructed cell types of the central brain, particularly the mushroom bodies and central complex, and not on truncated optic lobe cell types whose data are difficult to interpret (e.g. LC, LPC, LLPC, LPLC, and MC cell types in Fig. 8).

**Figure 8.**
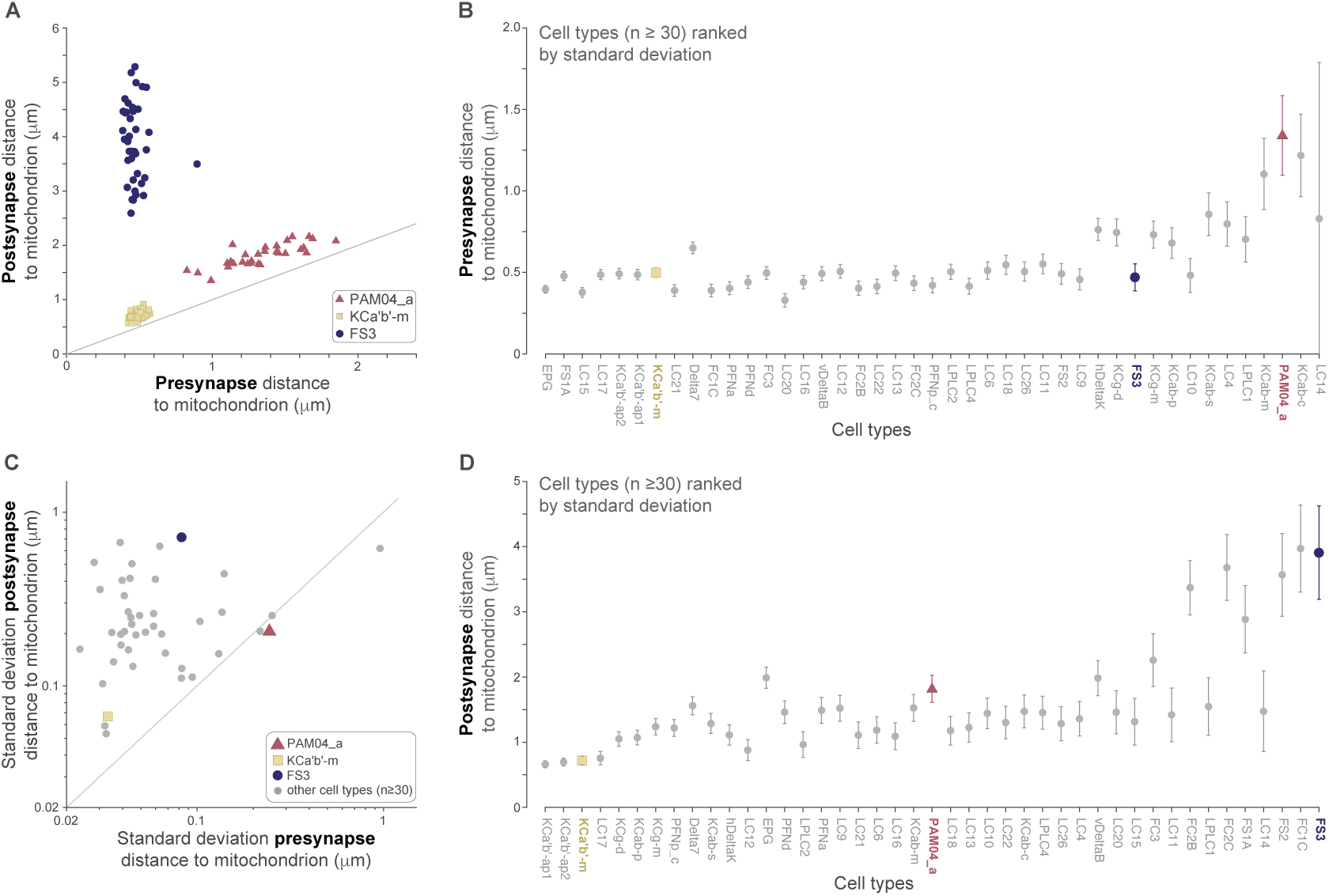
Mitochondrial placement near presynapses is more precise than at postsynapses. **A.** Examples of cell type variability in the distance to the nearest mitochondrion from presynapse (horizontal axis) and postsynapses (vertical axis). Median values shown for cells of select cell types with ≥ 30 cells each with >50 mitochondria: KCa’b’-m (n_cells_ = 119; yellow squares), PAM04_a cells (n_cells_ = 30; cherry triangles), and FS3 (n_cells_ = 40; blue circles); gray line indicates equal values. **B.** Mean and standard deviation of cell median distance to the nearest presynaptic mitochondria, for cell types with ≥ 30 cells, each with >50 mitochondria. In panels B-D, cell types from panel (A) are highlighted for reference. **C.** Standard deviation of the cell median distance to the nearest mitochondrion from presynapses (horizontal axis) and postsynapses (vertical axis), for cell types with ≥ 30 cells each with >50 mitochondria (gray circles), shown on logarithmic axes; gray line indicates equal values. **D.** Mean and standard deviation of cell median distance to the nearest postsynaptic mitochondria, for cell types with ≥ 30 cells, each with >50 mitochondria.

For KCa’b’ cell types, the distribution of distances is highly consistent (Fig. 8A; KCa’b’-m shown), indicating that the placement of their mitochondria is tightly regulated and important for synaptic function. The PAM04_a, KCab and KCg cell types have the unusual property that the distance to the nearest mitochondria is consistent at postsynapses, but highly variable between cells for presynapses (Fig. 8A; PAM04_a shown). The PAM04_a cells are involved in synaptic plasticity at their synapses in the mushroom bodies, and this spatial arrangement indicates that the cells may have differing degrees of involvement in synaptic plasticity. In contrast, cells of the fan-shaped body FS3 type are an extreme case of the typical pattern for most cell types, where the variability in the distance to the nearest mitochondrion is greater at postsynapses than at presynapses (Fig. 8A-D).

### Mitochondria and neurotransmitter type

In vertebrates, inhibitory cell types have been observed with mitochondria of a greater volume and number than in excitatory cell types (see ***Kann 2016*** for a review). However, it has not been possible to assess the generality of these findings across all cell types, due to a lack of access to large numbers of labeled cell types across different brain regions. Many of the cell types in our dataset have a known neurotransmitter expression, and for the remaining cell types we used the synapse-level neurotransmitter predictions of ***Eckstein et al. (2024)***, which were trained on connectome EM images and cell types whose neurotransmitter expression was confirmed with cell type-selective driver lines. For our analysis, we focused on the four most common and reliably predicted neurotransmitters by cell type (acetylcholine, dopamine, GABA, and glutamate; ***Eckstein et al. 2024***). In the *Drosophila* central nervous system, acetylcholine is the principal excitatory neurotransmitter, and glutamate is typically an inhibitory neurotransmitter (***Liu and Wilson, 2013***), unlike in vertebrates, although there are exceptions where, for example, different sets of downstream cells may express excitatory or inhibitory glutamate receptors (***Eddy et al., 2026***).

There are more than 1800 inhibitory cell types in our data, expressing GABA or glutamate. These cells have mitochondria that are much larger on average, by around 40%, than excitatory cells expressing acetylcholine (Fig. 9A; median mitochondrion volumes: Ach 0.122 µm^3^, GABA 0.164 µm^3^, Glu 0.168 µm^3^, Dop 0.112 µm^3^; all groups significantly different, *p* < 0.001, Kruskal-Wallis test with Bonferroni correction). Inhibitory cell types also have a greater density of mitochondria, as measured by the ratio of the total volume of mitochondria in a cell and the cell’s volume, greater by around 40% compared to cholinergic cell types (Fig. 9B; median mitochondrial volume densities: Ach 8.3%, GABA 11.7%, Glu 12.2%, Dop 9.4%; all groups significantly different, *p* < 0.001, Kruskal-Wallis test with Bonferroni correction). In contrast, the number of mitochondria per unit volume of the cell is similar, with a particularly wide distribution in dopaminergic cells (median mitochondrial number density: Ach 0.61 µm^-3^, GABA 0.62 µm^-3^, Glu 0.56 µm^-3^, Dop 0.75 µm^-3^). These findings establish the generality of greater mitochondrial investment in inhibitory neurons, compared to excitatory neurons, in the *Drosophila* central brain, as assessed across the more than 4300 cell types with known or predicted expression of these four neurotransmitters in our dataset.

**Figure 9.**
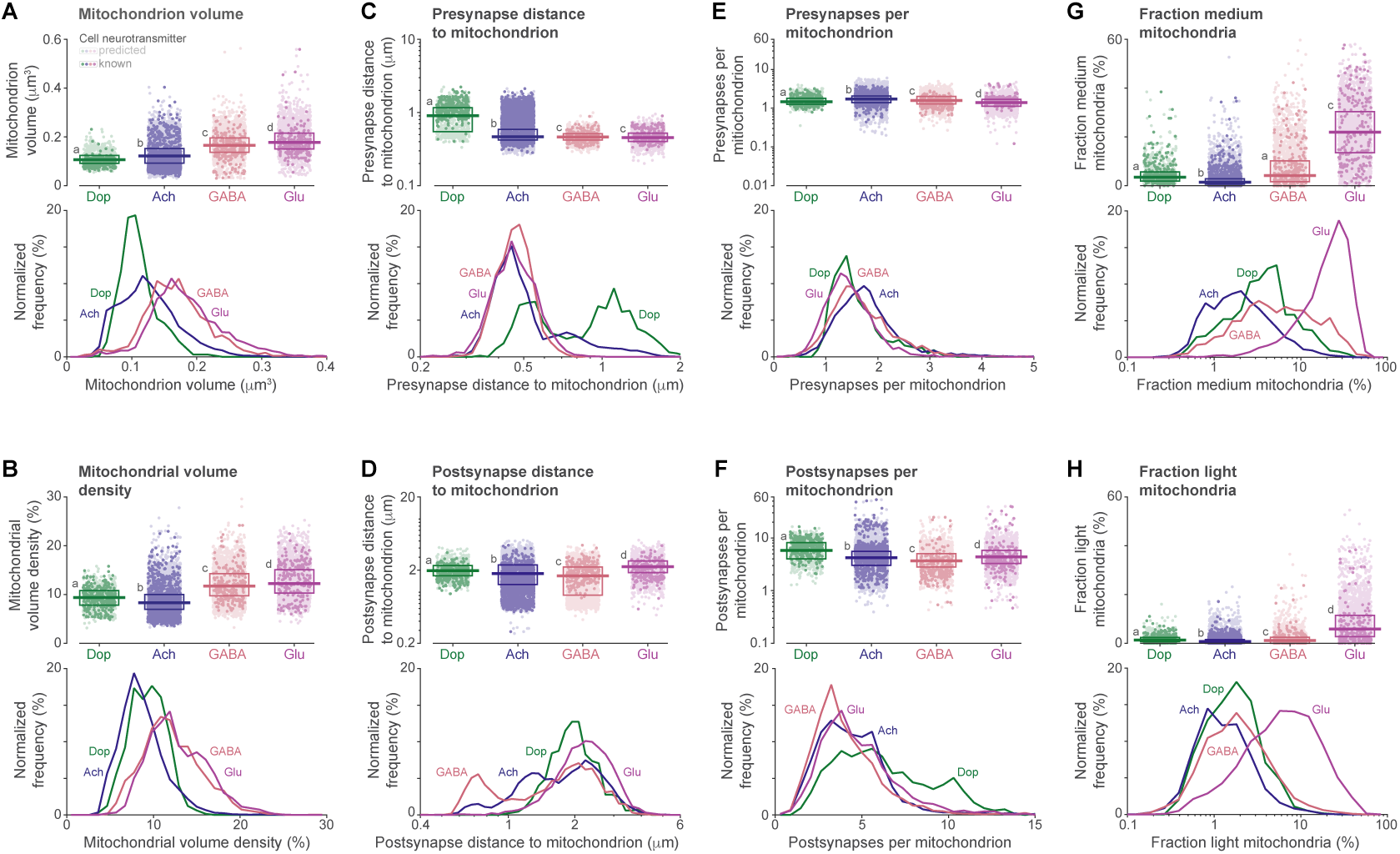
Mitochondria and neurotransmitters: greater investment in inhibitory neurons, distinct dopaminergic synapse placement, and lighter appearance in glutamatergic cells. Cells are grouped by neurotransmitter in all panels: in upper plots, dots indicate median values of individual cells of types with predicted (pale dots) or known (dark dots) neurotransmitter expression; box plots indicate the median and quartile ranges; letters (a,b,c,d) indicate significantly different distributions, assessed by Kruskal-Wallis test with Bonferroni correction for multiple comparisons; and lower plots show the normalized frequency distributions of cell median values. **A.** Mitochondrion volume: inhibitory (GABA and Glu) cells have substantially larger mitochondria than excitatory (Ach and Dop) cells. **B.** Mitochondrial volume density: the total volume of a cell’s mitochondria, divided by the cell’s volume. Inhibitory (GABA and Glu) cells have substantially higher densities of mitochondria than excitatory (Ach and Dop) cells. **C.** Presynapse distance to nearest mitochondria: dopaminergic cells have many more presynapses with mitochondria that are not nearby (i.e. ≥1µm) than other cell types. **D.** Postsynapse distance to nearest mitochondria: ells of all predicted neurotransmitter classes have wide distributions, without striking differences. **E.** Ratio of the number of a cell’s presynapses and mitochondria: the distributions are similar for all predicted neurotransmitter classes. **F.** Ratio of the number of a cell’s postsynapses and mitochondria: dopaminergic cells have a greater ratio than other cell types. **G,H.** Fraction of a cell’s mitochondria that have a medium (G) or light (H) appearance type: glutamatergic cells have a strikingly high fraction. Note that the fraction of dark mitochondria is redundantly equal to 100% minus the values in panels G and H, so is not plotted to avoid replotting the same data.

In vertebrates, dopaminergic cell types can vary in the dimensions of their synaptic mitochondria (***Chandra et al., 2019***), but how they vary in the distances between synapses and their nearest mitochondria is largely unknown. In our data, predicted and known dopaminergic cell types had a distinctive organization in the positioning of the nearest mitochondrion to a synapse (Fig. 9C-F). At presynapses, the median distance of the nearest mitochondrion was twice as far for dopaminergic cells, compared to the other neurotransmitter classes (Fig. 9C; median presynapse distance to mitochondrion: Ach 0.47µm, GABA 0.46µm, Glu 0.46µm, Dop 0.90µm; significant differences between Ach (a), GABA + Glu (b), and Dop (c) groups, *p* < 0.001, Kruskal-Wallis test with Bonferroni correction), and distances were greater in dopaminergic cells when only known cell types were analyzed (Fig. 9C, upper plots, dark dots; median values: Ach 0.72µm, GABA 0.45µm, Glu 0.49µm, Dop 1.11µm). In contrast, the distances to the nearest mitochondrion from postsynapses showed wide variability without such a striking difference for dopaminergic cells (Fig. 9D), and the number of presynapses per mitochondrion was also consistent across neurotransmitter classes (Fig. 9E). However, the number of postsynapses per mitochondrion is greater in dopaminergic cells (Fig. 9F; median postsynapses per mitochondrion: Ach 4.2, GABA 3.7, Glu 4.4, Dop 5.8; significant differences between all groups, *p* < 0.001, Kruskal-Wallis test with Bonferroni correction).

Cells predicted and known to express glutamate also had an unusually large fraction of mitochondria with medium (Fig. 9G; median values: Ach 1.4%, GABA 4.1%, Glu 21.9%, Dop 3.4%; significant differences between Dop + GABA (a), Ach (b), and Glu (c) groups, *p* < 0.001, Kruskal-Wallis test with Bonferroni correction) or light appearance types (Fig. 9H; median values: Ach 0.7%, GABA 1.2%, Glu 5.9%, Dop 1.3%; significant differences between all groups, *p* < 0.001, Kruskal-Wallis test with Bonferroni correction). Mitochondria play a key role in the biosynthesis of glutamate in neurons, and the more open cristae structure of mitochondria with medium and light appearance types may be related to this biosynthesis.

Together, the results of this section show that there are large, systematic differences in the properties of mitochondria of cells grouped by predicted and known neurotransmitter expression. Inhibitory cells have larger mitochondria, resulting in a greater mitochondrial density, while dopaminergic cells have mitochondria further from synapses and servicing greater numbers of postsynapses, and glutamatergic synapses have many more mitochondria with distinctively lighter appearances.

### Mitochondria at synapses with multiple postsynaptic partners

We hypothesized that presynapses with more postsynaptic partners would require a greater investment of mitochondria, as the release of more synaptic vesicles will require increased energy consumption (***Harris et al., 2012***). For example, mitochondria are positioned closer to larger synaptic clusters at synapses between astrocytes and neurons in mouse cortex (***Salmon et al., 2023***). In *Drosophila*, the majority of central brain synapses have multiple postsynaptic partners, and we refer to the number of postsynaptic partners as the synapse fan-out (Fig. 10). The postsynapses in our data are annotated by a single point, consistent with contemporary methods for synapse detection in insects (***Huang et al., 2018***; ***Buhmann et al., 2021***; ?), so we cannot readily measure how the total postsynapse area scales with the number of postsynaptic partners. Nevertheless, our analysis reveals how mitochondrial properties vary with fan-out on both sides of the synapse (Fig. 10).

**Figure 10.**
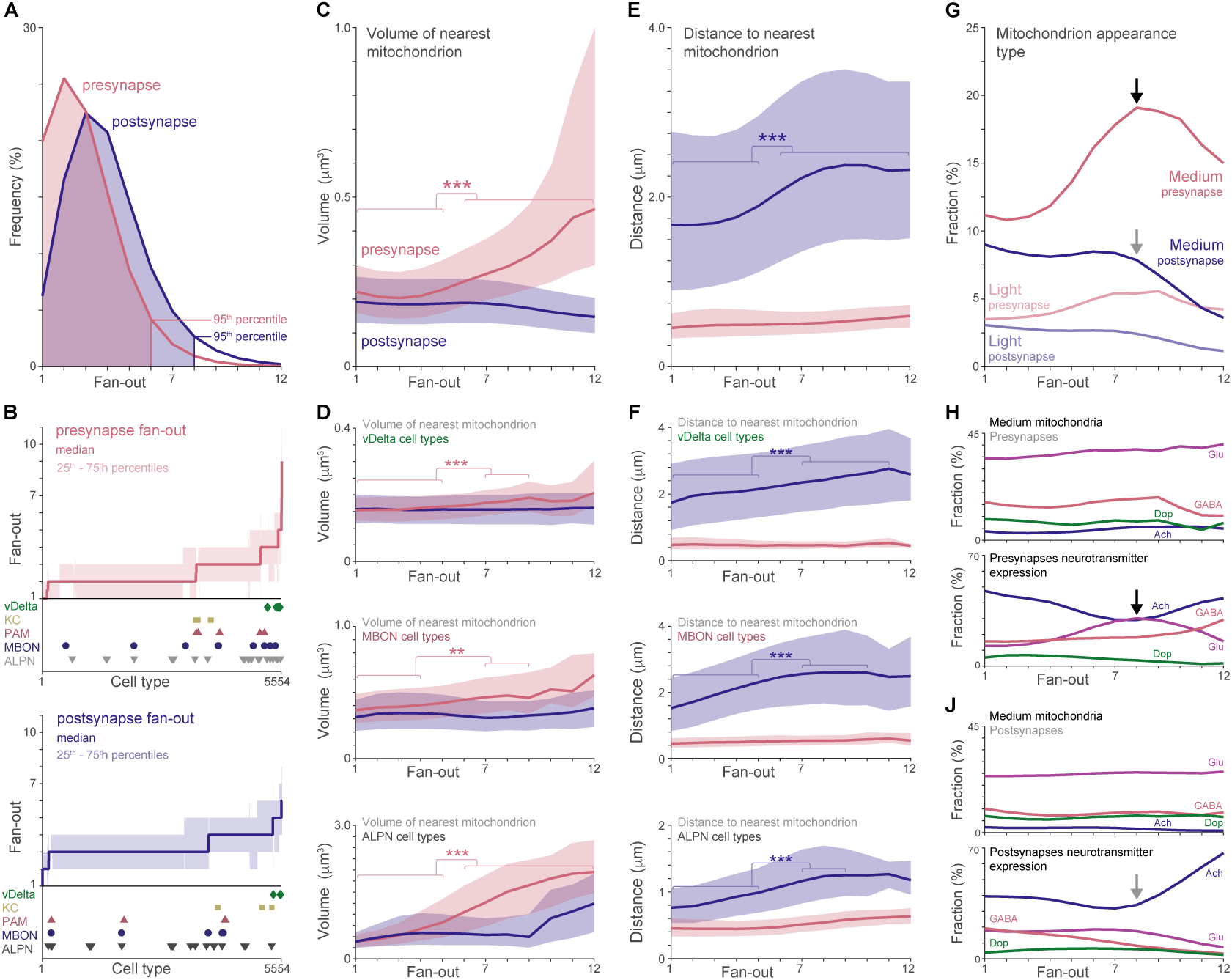
Larger presynaptic mitochondria and more distal postsynaptic mitochondria at synapses with multiple postsynaptic partners. **A.** Fan-out frequency. **B.** Cell types ranked by median synapse fanout for presynapses (top, cherry) and postsynapses (bottom, blue) with interquartile ranges (pale colors). Cell type medians annotated for: fan-shaped body vDelta cell types (green diamonds), KCs (yellow squares), PAMs (cherry triangles), MBONs (blue circles) and antennal lobe projection neuron (ALPN) cell types (gray triangles). **C,D.** Dependence on fan-out of median volume of the nearest mitochondrion to presynapses (cherry) and post-synapses (blue) for all cells (C) and for three groups of cell types (D) highlighted in panel B. Pale bands denote 33^rd^ and 67^th^ percentiles. **E,F.** Dependence on fan-out of distance to nearest mitochondrion for presynapses (cherry) and postsynapses (blue), for all cells (E) and for three groups of cell types (F) highlighted in panel B. Pale bands denote 33^rd^ and 67^th^ percentiles. **G.** Fraction of mitochondria appearance types for presynapses (cherry) and postsynapses (blue), for medium (full color) and light (pale color) appearance types. Arrows indicate fan-out values where the ratio of neurotransmitters expressed across the population of synapses affects fraction of medium mitochondria, highlighted in panels H and J. **H.** Top: for cells grouped by neurotransmitter expression, the fraction of medium mitochondria of presynapses does not markedly change with synapse fan-out. Bottom: the fraction of glutamatergic synapses is high at fan-out 8 (black arrow), corresponding to the peak in fraction of medium mitochondria in the presynapses of all cells (black arrow, panel G). **J.** As for panel H but for postsynapses. Gray arrow indicates where the fraction of synapses expressing acetylcholine increases, corresponding to the drop in panel G in the fraction of medium mitochondria at postsynapses of all cells (gray arrow in panel G). For all panels, asterisks denote significance: *** *p* < 0.001, ** *p* < 0.01, one-way ANOVA with Bonferroni correction.

The large majority (95%) of presynapses in our data have a fan-out between 1 and 6, and the large majority (95%) of postsynapses are partners in synapses with a fan-out between 1 and 8 (Fig. 10A). For individual cell types, the median fan-out is typically 2-4 for presynapses and 3-4 for postsynapses (Fig. 10B). Only rare cell types have high median fan-out, for example the vDelta cell types of the fan-shaped body brain region of the central complex (Fig. 10B; green diamonds). However, nearly all cell types have some synapses with higher fan-out, and more than 190,000 presynapses (3% of all presynapses in traced, uncropped named neurons) have a fan-out greater than 6.

Across all presynapses, the median volume of the nearest mitochondrion doubles as the fanout increases to 12 (Fig. 10C; *p* < 0.001, one-way ANOVA with Bonferroni correction; median volumes: 0.22 µm^3^ fan-out 1, 0.46 µm^3^ fan-out 12), while the volume of the nearest mitochondrion to a postsynapse decreases slightly (Fig. 10C; median volumes: 0.19 µm^3^ fan-out 1; 0.15 µm^3^ fanout 12). The trend for the volume of the nearest mitochondrion to a presynapse to increase with fan-out is consistent across groups of cell types, but varies widely in magnitude from, for example, +34% for vDelta cell types, to a substantial +400% for antennal lobe projection neuron cell types (Fig. 10D; median volumes: vDelta 0.154 µm^3^ fan-out 1; 0.206 µm^3^ fan-out 12; ALPN 0.389 µm^3^ fan-out 1, 1.96 µm^3^ fan-out 12).

In contrast to the greater investment in presynaptic mitochondria at synapses with a high fanout, the distance of the nearest postsynaptic mitochondrion increases with fan-out, which may reflect a reduced energy demand, a lower need for calcium regulation, or simply a lack of space at a crowded junction (Fig. 10E; *p* < 0.001, one-way ANOVA with Bonferroni correction). This trend is consistent across groups of cell types, but the range of distances varies widely (Fig. 10F; median postsynapse distances: 1.7-2.8 µm vDelta; 0.8-1.3 µm ALPN), accounting for the large variability around the median for the postsynapses of all cell types (Fig. 10E).

Finally, the fraction of mitochondria with medium and light appearance types doubles as the fan-out increases to 8 when calculated over all presynapses (Fig. 10G, black arrow). However, this effect is generated by a change in the ratio of synapses expressing different transmitters. Within each group of cell types expressing the same neurotransmitter, the fraction of medium mitochondria varies little with fan-out, and glutamatergic cell types have a high fraction, as expected (Fig. 10H, top). Meanwhile, the fraction of synapses expressing glutamate is high for a fan-out of 8 (Fig. 10H, bottom, black arrow). Similarly, the fraction of medium and light mitochondria at postsynapses decreases for postsynapses with a fan-out greater than 8 (Fig. 10G, gray arrow), because cholinergic synapses, which have a low fraction of medium and light mitochondria, dominate (Fig. 10J, gray arrow). So at synapses with high fan-out, glutamatergic presynapses dominate for fanouts of 8 or so, and cholinergic postsynapses are highly represented for fan-outs greater than 8, and this pattern accounts for variations in the mitochondrial appearance as the number of postsynaptic partners increases.

Overall, the results of this section indicate that mitochondria are larger at presynapses with more postsynaptic partners, reflecting a greater mitochondrial investment at synapses that are potentially releasing more synaptic vesicles (Fig. 10C). Meanwhile, the mitochondrial investment at postsynapses reduces with the number of postsynaptic partners, with mitochondria similarly sized and further away (Fig. 10E). For both of these effects, the magnitude is strongly affected by the cell type identity (Fig. 10D,F).

### Making mitochondria accessible for analysis

To facilitate the integration of mitochondrial data with neuronal connectome data, we have extended our public web interface, neuPrint (***Plaza et al. 2022***; https://neuprint.janelia.org). In neuPrint, neurons and synapses are recorded in a data model where they can be queried as a graph of connections and neighbors, and users can explore connectivity at varying levels of abstraction. For example, users can query the connections between brain regions, and then examine connections between specific pairs of neurons or the location of individual synapses. We have extended neuPrint to handle subcellular machinery and their interactions, and we describe the data model and further details in the Methods section. For each mitochondrion, we provide the location, appearance type, and shape information—the length of the axes of an ellipse fit to its volume. We also provide optional ‘CloseTo’ relations which encode the distance to nearby elements. As an example of the functionality that is now possible in neuPrint, investigators can use the Python interface to show the number of synapses within a certain distance of a mitochondrion.

## Discussion

We have created a dataset of annotated mitochondria in neural tissue that has an unprecedented number of mitochondria, synapses, neurons and cell types (Fig. 1). Our data quantifies the previously unknown ranges of mitochondrial properties in *Drosophila* neurons, enabling the identification of cell types with unusual values, including cells with pivotal roles in learning and memory (Fig. 2). The results demonstrate substantial differences in average mitochondrial properties of brain regions, and a much greater variability among cell types (Fig. 3). Cell types and groups of cell types have distinctive patterns of synaptic placement (Figs. 4,5), typically with a mitochondrion within one micron of presynapses (Fig. 6). Mitochondria are larger and closer at presynapses than at postsynapses (Fig. 7), and the placement of mitochondria is particularly precise in cell types that play a key role in sensory learning, Kenyon cells (Fig. 8). We have confirmed, using thousands of cell types in our sample, that inhibitory interneurons are more densely packed with mitochondria than excitatory neurons, that dopaminergic neurons have distinct synaptic mitochondrial positioning, with mitochondria more distant at presynapses, and that glutamatergic neurons have a much greater fraction of mitochondria with a light appearance (Fig. 9). We have also identified organizational principles for mitochondria nearest synapses with multiple postsynaptic partners: as the number of partners increases, the nearest presynaptic mitochondrion is larger while the mitochondria nearest the postsynaptic contact sites are further away (Fig. 10). Finally, we have extended our publicly available web interface, neuPrint, to support the co-investigation of mitochondria, synapses, neurons, cell types and brain regions across this volume that comprises the majority of the central brain of *Drosophila*.

Despite the comprehensive reach of our dataset across brain region and cell types, there are limitations to our data. Some neurons are incomplete, particularly cell types with neurites in brain regions outside the sample, including ascending and descending neurons of the ventral nerve cord, and neurons extending into the brain regions processing visual information, the optic lobes (***Nern et al., 2025***). Nevertheless, even in the optic lobes, subsequent researchers have been able to analyze our data to identify properties of mitochondrial organization in visual projection neurons, cells connecting the optic lobe with the central brain, including how mitochondrial morphology varies within and between cell types, how mitochondria vary between axons and dendrites, and how the placement of mitochondria can vary by local region and the organization of postsynaptic partners outside the optic lobes, for example, in Kenyon cells (***Sager et al., 2026***). Our analysis is also a snapshot of the state of mitochondria in the brain of a young, naive fly fed *ad libitum*. Environmental experience, neuronal activity, physiological state, sex, age, stress and nutrition all have profound effects on mitochondria (***Uytiepo et al., 2025***; ***Cserép et al., 2018***; ***Sarnataro et al., 2025***; ***Kristensen et al., 2019***; ***Li et al., 2024***; ***Hamer et al., 2025***; ***Kuszak et al., 2018***), so these and other factors may alter the mitochondrial properties in neurons: our work provides a crucial starting point for further studies. Finally, the statistical variation of mitochondrial characteristics, and the strong dependence on cell type, mean that detailed conclusions for a given cell type will likely require analyzing many cells of the same cell type. In the Hemibrain sample many cell types have only a few instances, and the application of our methods to the optic lobe of recent whole brain connectomes (***Berg et al., 2025***; ***Schlegel et al., 2024***), where there are many cell types with hundreds of instances each, and cell types have been reconstructed to the greatest level of completion of large-scale *Drosophila* connectomes thus far (***Nern et al., 2025***), will overcome this limitation.

Our results are consistent with many studies that have focused on specific brain regions or cell types, and they significantly develop those findings by providing a quantitative context of the majority of the central brain within which the findings can be appreciated. Here, we briefly focus on two such results. First, many studies have observed that mitochondria are usually present at presynapses, but are often not present at postsynapses, in excitatory neurons of wide-ranging tissue from mouse cortex (***Kasthuri et al., 2015***; ***Turner et al., 2022***) to the suprachiasmatic nucleus (***Calligaro et al., 2023***); it was surprising, therefore, that relatively few presynapses of hippocampal pyramidal cells, which play a vital role in energetically costly short-term memory, have a nearby mitochondrion (***Shepherd and Harris, 1998***; ***Smith et al., 2016***; ***Bloss et al., 2018***). In our data, most cell types across brain regions have mitochondria near almost all their presynapses, but specific cells involved in learning and memory do not, the KC and PAM cell types, and the unusual extent of this organization is precisely enumerated (Fig. 6B). These results may indicate that a lack of a mitochondrion at a presynapse may facilitate synaptic plasticity, for example by altering presynaptic calcium levels (***Kwon et al., 2016***; ***Vaccaro et al., 2017***; ***Li et al., 2020***), or simply that when a neuron has a set of synapses whose strength can be altered, there will be some synapses that remain quiet and do not require resources (***Levy and Baxter, 1996***).

Second, previous work has identified that inhibitory neurons often have high levels of mitochondria (***Kageyama and Wong-Riley, 1982***, ***1984***; ***Shepherd and Harris, 1998***; ***Kann, 2016***; ***Turner et al., 2022***), especially in interneurons capable of sustaining high levels of activity (***Cserép et al., 2018***; ***Kann, 2016***). The generality of these observations was unclear, however, as these studies have focused on specific brain regions or cell types. Our results show that across thousand of cell types and many brain regions, inhibitory cell types have around 40% greater mitochondrial investment than cells expressing acetylcholine, the predominant excitatory neurotransmitter in the fly central brain, and they quantify the fraction of cell types that may nevertheless oppose this trend when assessed in isolated pairs (Fig. 9B). What are the principles driving mitochondrial investment in inhibitory neurons? One function of inhibition is to act as a brake on the activity of excitatory cells: for example the inhibitory APL cell (Fig. 7A) regulates the activity of mushroom body cells including KCs (***Amin et al., 2020***), and has a high mitochondrial volume density (median vol. density: 14%; 83^rd^ percentile). Accordingly, one component of this mitochondrial investment may be to save metabolic resources across the brain through greater deployment in cells that reduce the activity of others. Inhibitory neurons are also employed in circuits requiring sustained high activity levels, for example generating sustained oscillatory activity in the hippocampus (***Kann, 2016***). In the fly, inhibitory ring neurons, for example, are key components of the circuitry supporting the sustained activity of direction-encoding neurons (***Seelig and Jayaraman, 2013***; ***Turner-Evans et al., 2020***), and ring neurons have among the highest mitochondrial density of all cell types. For example, ER2_b neurons encode polarized light signals supporting the sense of direction (***Hardcastle et al., 2021***) and their mitochondrial volume density is in the 96^th^ percentile (median vol. density: 18%). Thus, tonic or sustained activity may be a further, complementary component of mitochondrial investment in inhibitory neurons.

Two further functional explanations for the investment of mitochondria in a cell type can be explored in our data: speed and synaptic plasticity. In sensory neurons, the energetic cost of encoding changes in the sensory input increases dramatically with the speed of changes in the encoded sensory signals, a cost that arises from the need to use ATP to maintain the cell’s ionic fluxes (***Niven et al., 2007***). From these observations, it is natural to infer that rapid neural processing may carry a higher energetic cost and require a correspondingly greater investment of mitochondria. In our data, neurons along the visual escape pathways have low mitochondrial densities, indicating that rapid processing does not necessitate high mitochondrial investment (e.g. LC4, Giant Fiber, LPLC2 ***Ache et al. 2019***; Suppl. Data Tables 1,2). Cells along the visual escape pathway have rapidly adapting responses to sensory stimulation, which may allow them to rapidly encode the onset of sensory stimuli without great energetic cost (***Benda, 2021***). For synaptic plasticity in flies, since learning can be vulnerable to starvation and depends on the metabolic state (***Plaçais and Preat, 2013***; ***Plaçais et al., 2017***; ***Amrapali et al., 2026***), it is also natural to infer that cells expressing synaptic plasticity require a greater investment of mitochondria. Our analysis of memory-supporting Kenyon cells indicate that while these cells have many specializations in the organization of their mitochondria, including precise localization at presynapses (Fig. 8), the cells’ synaptic plasticity does not necessitate high mitochondrial investment, consistent with the importance of glia for supplying the cells’ energetic requirements (***Rabah et al., 2023***).

EM reconstructions provide the gold standard for quantifying the organization of mitochondria within and between neurons. Our data demonstrate how large scale connectomes allow this organization to be analyzed across brain regions and across a comprehensively large sample of cell types. The results show how this approach can be used to appreciate the specificity of mitochondrial organization in certain cell types, notably in memory-supporting Kenyon cells, and identify potential principles for varied mitochondrial investment across the whole brain.

## Methods

### EM volume

We analyzed images of the Hemibrain sample that were taken with a scanning electron microscope and focused ion beam milling between sections (***Scheffer et al., 2020***). The images have an isotropic resolution of 8 nm per voxel, a level of spatial detail that enables detailed reconstructions of mitochondria (Fig. 1). However, the dataset was optimized for purposes that affect mitochondrial segmentation. First, the sample fixation and staining were optimized to identify cell membranes and synapses, not mitochondria. The hemibrain was prepared with the PLT-LTS method, which involves dehydration by progressive lowering of temperature, along with low temperature staining (***Lu et al., 2022***). This method helps to provide uniform osmication and staining, with less chance of distorting the fine structure. Second, the samples were optimized for reconstructing the cells of the mushroom body and central complex brain regions; in other brain regions, specifically the truncated optic lobes, the contrast varies and many mitochondria are stained black with little internal structure, affecting the segmentation. Despite these issues, the mitochondria are readily identifiable (Fig. 1), and we detail the accuracy of our mitochondrial segmentation below.

### Mitochondria segmentation

In our prior work, mitochondria masks were created for the Hemibrain dataset to prevent mitochondria from interfering with the segmentation of neurons (***Scheffer et al., 2020***). This prior goal did not require precise masks and created mitochondrial segmentation artifacts: for example, single mitochondria were segmented into fragments of several classes, such as small cells as well as mitochondria. Post-processing was therefore needed to turn the mitochondria masks into mitochondria objects. For this reason, our segmentation methods comprise an augmented version of the segmentation previously used for cells (***Scheffer et al., 2020***) followed by manual correction steps, and are not an automated pipeline for the detection of mitochondria from EM images.

By manual inspection, we determined that the mitochondria in the sample varied in the intensity of their staining, and that including this information improved the performance of the automatic segmentation. Therefore, we included three categories of mitochondrial appearance type in the creation of the manually labeled training data (Fig. 1). We manually created ground truth data for training the automatic segmentation of mitochondria from the images: two proofreaders annotated all the mitochondria within small cubes taken from each region of interest. In total, we annotated 13,000 mitochondria, annotating 739,000 voxels of dark mitochondria, 537,000 voxels of medium mitochondria, 473,000 voxels of light mitochondria, and 34,000,000 voxels of background that contained no mitochondria.

We report our mitochondria appearance type directly, after exploring the application of existing terms to our data. The seminal work of ***Hackenbrock (1966)*** established two main configurations in mammalian tissue: ‘orthodox’ and ‘condensed’ (see, for example, his Fig. 9 for orthodox and Fig. 3 for condensed). We observed no mitochondria that look like his ‘condensed’ configuration, and our ‘light’ mitochondria match those he calls ‘orthodox’, as is also shown in Fig. 1 of ***Scalettar et al. (1991)***. Conversely, ***Frey and Mannella (2000)***, referring to Hackenbrock, assert that “mitochondria observed in situ are almost always found in the ‘orthodox’ conformation”. According to this, our dark mitochondria would be their orthodox configuration. Because of these ambiguities, we report our mitochondria appearance type as observed.

Next, mitochondria masks were computed at Google, using the ground truth annotations to train the same model architecture that was used for the previous 7-class segmentation of the Hemi-brain sample (***Scheffer et al., 2020***). The differences were that for the mitochondrial segmentation, 4 classes were used (background and 3 mitochondria appearance types), and the training examples were randomly rotated in three dimensions. The results of this step were three categories of mitochondria masks containing voxels of different appearance types: dark, medium, and light, respectively.

Extracting instances of mitochondria from the masks required combining adjacent mask areas. Many mitochondria were initially segmented into fragments of several classes, for example dark and medium masks. Consistent with our observations of mitochondria composed of several classes, mitochondria with mixed configurations (orthodox and non-orthodox) have also been observed (***Smith et al., 2016***). When two mitochondria masks of different appearance types touched, we merged them and assigned the appearance type with the most voxels.

Extracting instances of mitochondria also required processing artifactual segments of mitochondria as small neurons. In the previous version of the segmentation (***Scheffer et al., 2020***), some mitochondria were over-segmented as their own objects, rather than being properly merged into their surrounding neurons. This had no effect on neural connectivity, but resulted in mitochondria being classified as neurons. To determine which neurons such mitochondria fragments should be merged to, we used the following two-step procedure. First, the convex hulls of adjacent neurons were computed in a neighborhood around the over-segmented mitochondrion. Second, each fragment was then merged into the neuron whose convex hull overlapped with that fragment more than any other neuron’s convex hull. This procedure was effective in handling a common error, where two mitochondria from different cells are separated by the narrow, dark cell membranes; other simpler procedures, such as assigning fragments according to the proportion of direct adjacencies, or filling holes using watershed algorithms, failed to handle this error.

### Accuracy of mitochondria segmentation

We characterized the precision (fraction of detections which are true positives) and recall (fraction of detections relative to all mitochondria) of our mitochondrial segmentation as follows.

To quantify the precision of the mitochondrial segmentation, proofreaders examined a random selection of mitochondria identified by our segmentation. For each selected mitochondrion, the proofreader reported whether it was indeed a mitochondrion and, if so, selected a point within the mitochondrion in the EM images. We then checked whether the point was inside any mitochondria mask, and whether the point was inside the randomly selected mitochondrion. In 99% of cases, the proofreader’s point was within a mitochondrion mask. In 85% of cases, the proofreader’s point was in the mask of the randomly chosen mitochondrion instance. The lower accuracy for this second measure was primarily due to instances where a mitochondrion was segmented into two or more mitochondria close together. These errors were difficult to eliminate, because mitochondrial fission and fusion events are common in vivo: in fission, one mitochondrion can split into two mitochondria; and in fusion, two mitochondria can unite to form one mitochondrion. Across such a large sample volume, the sample contains many mitochondria that have just initiated or completed fission or fusion, events that are easy to identify manually but harder to classify automatically. This class of error creates a small bias in our results towards smaller mitochondrial volumes and greater mitochondrial numbers per cell, but has little effect on the total mitochondrial volume per cell and on our analysis of the distance of mitochondria closest to synapses.

The recall of the mitochondrial segmentation near synapses was particularly important for the accuracy of our analysis of the mitochondria closest to synapses. To quantify this recall, proofreaders examined a random selection of synapses. The proofreader examined a 1 µm spherical volume surrounding each selected synapse, and selected the nearest point within the mitochondrion closest to the synapse within the cell, if there was one present in the spherical volume. We then checked whether this point was indeed marked as a mitochondrion within our data. We performed this analysis on the central brain regions, including the mushroom body and fan-shaped body, where cells are reconstructed to a higher level of completion than other brain regions, and not on the optic lobes: the measure was >95% across all these brain regions.

### Synapse-to-mitochondrion distances

To calculate the distance of the nearest mitochondrion from a synapse, we used traced cells in the Hemibrain dataset v1.2.1 that are not cropped (in neuPrint: status=’Traced’, cropped=’False’). In this dataset, incompletely traced neurons are listed as fragments and were not analyzed. For all synaptic elements, presynapses and postsynapses, the locations are represented by single points. To briefly describe the synapse identification process, the centers of T-bars and PSDs were manually labeled in a training dataset, which was used to train a machine learning model to predict the centers of T-bars and PSDs throughout the volume, following an iterative approach where model-retraining was interleaved with manual proofreading (for full details, see ***Scheffer et al. 2020***).

We computed the distance as the path length within the neuron from the synapse to the nearest edge of the nearest mitochondrion. This was an intensive computation, so to reduce the required number of calculations we performed it at a resolution of 64 nm, and not the full 8 nm resolution of the images. This resolution gives a high degree of accuracy for distances that are typically of the order of 1000 nm. The distances are expressed, however, in voxel units, where the distance of one voxel unit is 8 nm. The computation was able to allow for small gaps in the cell segmentation, but not large gaps: small gaps were traversed in 0.85% of synapse-to-mitochondria paths, and large gaps were less frequent. At the reduced resolution used in the worst cases, the accuracy of the distance calculation was ±28 voxels, equivalent to 0.22 µm; for comparison, the median mitochondrion length is 1 µm.

For 0.96% of all synapses in traced cells, no path to a nearby mitochondrion could be found. We did not find a nearest mitochondrion when the synapse was on a neural fragment isolated from its nearest mitochondria. We also introduced a distance threshold: when there was no mitochondrion within 10 µm, we determined that there was no nearest mitochondrion.

### Mitochondrial morphology

The shape of each mitochondrion was estimated by fitting an ellipsoid to the connected voxels. This yields shape descriptors describing the lengths of the three axes of the ellipsoid, *R*_1_, *R*_2_ and *R*_3_. We have used *R*_1_ in our current analysis, and not *R*_2_ and *R*_3_, but we have recorded all three parameters in our public web interface, neuPrint, which we describe next.

### neuPrint

The organization of data in neuPrint is shown in Figure 11. A principal extension is the ability to declare arbitrary elements (for example, synapses and mitochondria are elements) that have two key properties: a location and type. A second extension is that non-synapse elements are grouped within a node type called an ElementSet. We have grouped the mitochondria for each neuron within an ElementSet which then points to the specific mitochondria. Summary information about the number or distribution of mitochondria are provided at both the ElementSet and brain region level.

**Figure 11.**
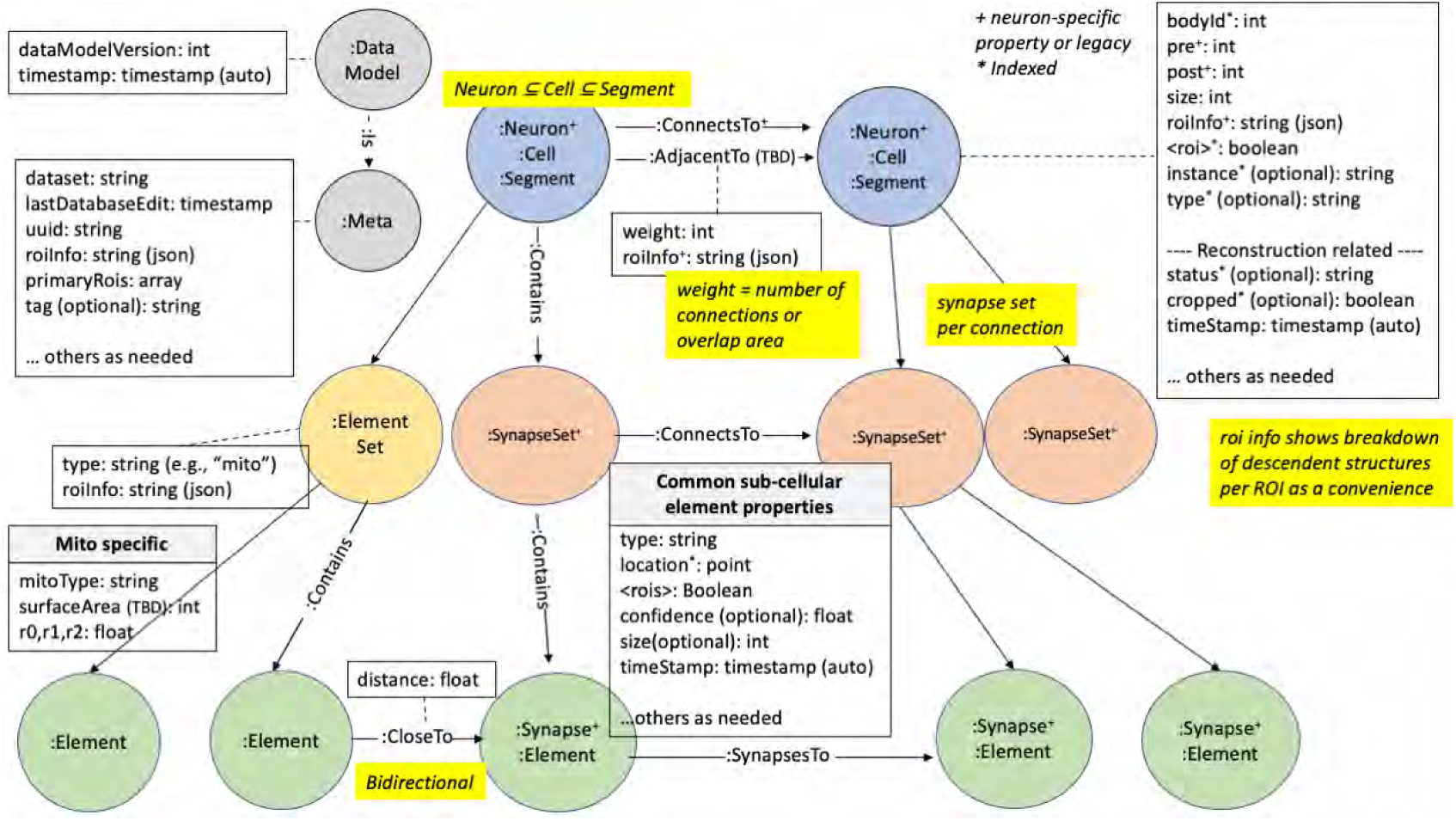
Organizing schema for neuPrint. Our extension to neuPrint supports the inclusion of sub-cellular components, specifically mitochondria for the Hemibrain dataset.

The nearest mitochondrion to each synapse is recorded in neuPrint via a: CloseTo relationship for the synapse, with distance stored as a property of the relationship. For synapses with no closest mitochondrion, we did not record a: CloseTo relationship. While the distance encoded in the ‘CloseTo’ property is the shortest path within the containing cell that connects the two elements, the investigator also can calculate other distances in neuPrint, such as the Euclidean distance.

In neuPrint, we have maintained backwards compatibility for neuPrint, and created the possibility of nested hierarchies of structures, such as sub-organelle morphologies where image resolution permits. The data model is also compatible with non-neuronal tissue: for example, proximity between cells could be recorded instead of the number of synapse connections. We have enhanced the neuPrint web and Python interfaces to include these additions to the data model. The web application can show distributions of mitochondria, with breakdowns per mitochondria appearance type per brain region for each neuron. A user can also view low-level connections between elements by specifying location, or by clicking through a given neuron and its element set. Both web and Python interfaces provide a general spatial query function to find elements of a specified type or types near a given coordinate. Finally, we note that in future updates of the Hemibrain dataset, some aspects of the data model related to mitochondria that may change, for example the spelling of cell type names.

### Neurotransmitter predictions

To obtain the neurotransmitter expressed at a synapse, we relied on the predictions from ***Eckstein et al. (2024)***. From those, we assigned the primary neurotransmitter of a cell as the most common highest predicted probability over all presynapses of that neuron, for cells with >100 presynapses. We then assigned a neurotransmitter to a cell type when the predictions agreed for ≥80% cells of that type; for cell types not meeting this consensus, the cell type neurotransmitter prediction is ‘unknown’. Our analysis concentrates on the four most common and reliably predicted neurotransmitters by cell type (acetylcholine, dopamine, GABA, and glutamate). For cell types where the expressed neurotransmitter is known, we used that neurotransmitter expression for all cells of that cell type. For example, we labeled the Kenyon cells as expressing acetylcholine (***Barnstedt et al., 2016***), rather than as dopaminergic, as they are labeled in ***Eckstein et al. (2024)***.

## Data analysis

### Code Availability

Code for analyzing the data is written in python and available as python notebooks, available at github.com/kitlongden/mitohb_2026. The analysis code is written using functions documented in the neuprint-python application programming interface (connectome-neuprint.github.io/ neuprintpython/docs). These functions implement common search queries to the database in which the data is stored. Statistics and multi-panel figures are generated using MATLAB (version 25.2.0, R2025b; The MathWorks Inc. Natick, MA, USA), with code also available at github.com/kitlongden/mitohb_2026. Neuroglancer (github.com/google/neuroglancer) is used to visualize mitochondria in segmented cells (Figs. 1D, 2A, 4, 7A,E).

### Cell and ROI volumes

Cell volume is stored in neuPrint as a property of the cell, and these stored values were used to calculate the mitochondrial number density and mitochondrial volume density (Fig. 2E,G, 9B). Calculating the volume of part of a cell from cell skeleton information requires consideration, because cell skeletons are created and stored at a spatial resolution that varies across cells: large cells with complex arbors having a coarser skeleton spatial resolution than small neurons. As a result, skeletons may need to be recomputed at the same spatial scale to make comparisons between, for example, the axonal or dendritic arbors of different cell types. ROI volumes are calculated using the natverse toolbox for combining and analyzing neuroanatomical data (***Bates et al., 2020***).

### Synapse fan-out

The synapse fan-out of presynapses is calculated by first taking all traced and uncropped cell types, and identifying all of their paired synaptic connections. These synaptic connections are grouped by presynapse to calculate, for every presynapse, the number of postsynaptic partners, which is the presynapse fan-out. These values are then stored in a look up table that includes the identity of every presynapse and postsynapse and provides a fan-out value for every synaptic connection. From this look up table, the postsynapse fan-out is readily identified. For synapses with very large fan-outs, ≳20, these may include synapses with multiple release sites very near to each other, that have been annotated with single point, and manual inspection is recommended.

## Author contributions

SB, MJ, EN, CO, and SP created the neuron segmentation; MJ, CO, and EN created the mitochondria semantic segmentation; SB created the mitochondria instance segmentation and calculated synapse-to-mitochondria distances; KL, CO, PR, and LS analyzed mitochondria-synapse relationships; KL and LS analyzed mitochondria-neurotransmitter relationships; SB, EN, CO, EP, PR, NS, ST, CW, and EY did the proofreading; SB, KL, EN, CO, SP, PR, and LU did quality control; SB, JC, MJ, KL, SP, and LU contributed software; KL, SB, MJ, PR, and LS wrote the paper; KL revised the paper, in coordination with LS, PR, MD, TM and MR; SB, EN, SP, and PR coordinated internal efforts; PR coordinated annotation; SB and SP managed the overall effort.

## Acknowledgements

Thanks to members of the Drosophila Connectomics Group, Jefferis lab, Devine lab, Reiser lab, Prieto-Godino lab, and Prof. Joe Bateman for comments on the revised manuscript. This work is funded by the Howard Hughes Medical Institute through its support of the Janelia Research Campus and the FlyEM Team Project. MJ was supported by Google Research. Proofreading of the Hemibrain dataset was also supported by a Wellcome Trust Collaborative Award (203261/Z/16/Z). PR was additionally supported by the National Institute of Mental Health of the National Institutes of Health under Award Number R24MH114785. The content is solely the responsibility of the authors and does not necessarily represent the official views of the National Institutes of Health. TFM and MJD were supported by The Francis Crick Institute, which receives its core funding from Cancer Research UK (CC2206), the UK Medical Research Council (CC2206), and the Wellcome Trust (CC2206). This article is subject to HHMI’s Open Access to Publications policy. HHMI lab heads have previously granted a nonexclusive CC BY 4.0 license to the public and a sublicensable license to HHMI in their research articles. Pursuant to those licenses, the author-accepted manuscript of this article can be made freely available under a CC BY 4.0 license immediately upon publication.

## Supplemental Data

Supplementary Data Table 1: Mitochondrial properties of cells.

Supplementary Data Table 2: Mitochondrial properties of cell types.

